# ESCRT machinery mediates selective microautophagy of endoplasmic reticulum

**DOI:** 10.1101/661306

**Authors:** Jasmin A. Schäfer, Julia P. Schessner, Peter W. Bircham, Takuma Tsuji, Charlotta Funaya, Katharina Schaeff, Giulia Ruffini, Dimitrios Papagiannidis, Michael Knop, Toyoshi Fujimoto, Sebastian Schuck

## Abstract

ER-phagy, the selective autophagy of endoplasmic reticulum (ER), safeguards organelle homeostasis by eliminating misfolded proteins and regulating ER size. ER-phagy can occur by macroautophagic and microautophagic mechanisms. While dedicated machinery for macro-ER-phagy has been discovered, the molecules and mechanisms mediating micro-ER-phagy remain unknown. Here, we first show that micro-ER-phagy in yeast involves the conversion of stacked cisternal ER into multilamellar ER whorls during microautophagic uptake into lysosomes. Second, we identify the conserved Nem1-Spo7 phosphatase complex and ESCRT proteins as key components for micro-ER-phagy. Third, we demonstrate that macro- and micro-ER-phagy are parallel pathways with distinct molecular requirements. Finally, we provide evidence that ESCRT proteins directly function in scission of the lysosomal membrane to complete the microautophagic uptake of ER. These findings establish a framework for a mechanistic understanding of micro-ER-phagy and, thus, a comprehensive appreciation of the role of autophagy in ER homeostasis.

## INTRODUCTION

Cells profoundly remodel their organelles in response to changing physiological demands. The architecture of the ER in particular undergoes extensive changes during differentiation, stress and disease. These changes include massive ER expansion when cells boost lipid synthesis or protein secretion (Pathak et al., 1986; Wiest et al., 1990). ER expansion can be reversed by organelle-selective autophagy (Bohlender and Weibel, 1973; Fumagalli et al., 2016). In addition, misfolded proteins in the ER can be eliminated by autophagy (Ishida et al., 2009; Forrester et al., 2019). Hence, autophagy promotes ER homeostasis by regulating organelle size and mediating protein quality control.

Two types of autophagy appear to be universally conserved in eukaryotes: macro- and microautophagy (Galluzzi et al., 2017). During macroautophagy, autophagic cargo is enclosed in autophagosomes, which then fuse with endolysosomes. During microautophagy, autophagic cargo is directly engulfed and taken up by endolysosomes. Both macro- and microautophagy can selectively target certain organelles or operate non-selectively, giving rise to a variety of autophagy pathways.

Macroautophagy requires the core autophagy machinery, a conserved set of proteins essential for autophagosome formation (Mizushima et al., 2011; Farré and Subramani, 2016). Organelle-selective macroautophagy additionally involves autophagy receptors, which facilitate inclusion of target organelles into autophagosomes. Many other proteins participate in macroautophagy (Galluzzi et al., 2017; Zhao and Zhang, 2019). Of note, ESCRT (endosomal sorting complexes required for transport) proteins are needed for efficient macroautophagy (Filimonenko et al., 2007). ESCRT proteins mediate membrane remodelling and scission in various processes, including multivesicular endosome formation and cytokinesis (Schöneberg et al., 2016; McCullough et al., 2018). In autophagy, ESCRT proteins have been proposed to facilitate autophagosome closure, which involves membrane budding away from the cytosol and is therefore topologically equivalent to other ESCRT-mediated processes (Hurley, 2015; Spitzer et al, 2015; Takahashi et al., 2018; Zhou et al., 2019).

Microautophagy is less well understood than macroautophagy. In yeast, the core autophagy machinery is required for microautophagy of peroxisomes and portions of the nucleus (Mukaiyama et al., 2002; Krick et al., 2008), may be required for microautophagy of lipid droplets under certain conditions (van Zutphen et al., 2014; Wang et al., 2014; Oku et al., 2017), is not required for microphagy of ER (Schuck et al., 2014), and may promote efficiency of microautophagy of cytosol (Müller et al., 2000; Oku et al., 2017). Several factors could contribute to this puzzling picture. First, the term ‘microautophagy’ likely conflates distinct types of autophagy (Schuck et al., 2014; Oku and Sakai, 2018). Second, microautophagy may in some settings indirectly depend on the core autophagy machinery. Microautophagy consumes endolysosomal membrane, which may need to be replenished by macroautophagy to maintain endolysosome function (Müller et al., 2000). Third, it remains possible that some microautophagy pathways utilize only part of the core autophagy machinery. Importantly, ESCRT proteins are required for many types of microautophagy (Sahu et al., 2011; Liu et al., 2015; Oku et al., 2017; Tsuji et al., 2017; Mejlvang et al., 2018). While the topological equivalence of microautophagy and other ESCRT-dependent processes is compelling, a direct involvement of ESCRT machinery in microautophagy has been difficult to demonstrate.

ER-phagy, the selective autophagy of ER, can occur by macroautophagy in both yeast and mammals, and several macro-ER-phagy receptors have been identified (Hamasaki et al., 2005; Khaminets et al., 2015; Mochida et al., 2015; Fumagalli et al., 2016; Grumati et al., 2017; Smith et al., 2018; An et al., 2019; Chino et al., 2019). Micro-ER-phagy has been found in yeast, in which protein folding stress in the ER induces spherical ER whorls that undergo microautophagy independently of the core autophagy machinery (Schuck et al., 2014). At present, the machinery for micro-ER-phagy is unknown, which hinders analysis of its mechanism, clarification of its physiological functions and assessment of its conservation in mammals.

Here, we define the dynamic membrane rearrangements during micro-ER-phagy in yeast, identify key components of the molecular machinery involved, show that macro- and micro-ER-phagy are genetically separable, parallel pathways and provide evidence that ESCRT proteins mediate lysosomal membrane scission during microautophagic uptake of ER.

## RESULTS

### Expression of ER-resident transmembrane proteins can induce micro-ER-phagy

Electron micrographs of yeast treated with the ER stressors DTT or tunicamycin revealed ER whorls as ring-shaped structures (Figure S1A; Schuck et al., 2014). To visualize ER whorls by light microscopy, we imaged yeast expressing the general ER marker Sec63-GFP. This approach highlighted pronounced expansion of the organelle during ER stress (Figure S1B; Schuck et al., 2009). However, neither Sec63-GFP nor other transmembrane or lumenal ER markers tagged with diverse fluorescent proteins labelled structures resembling whorls. A possible explanation could be that whorls contain low concentrations of certain ER proteins. To address this possibility, we applied freeze-fracture electron microscopy, which visualizes transmembrane proteins as intramembrane particles (Kuby and Wofsy, 1981). Freeze-fracture images revealed DTT-induced multilayered membrane spheres in the cytosol and the vacuole, the yeast lysosome (Figure 1A). Strikingly, these spheres consisted of smooth membranes largely devoid of intramembrane particles, in contrast to regular ER. Hence, ER whorls induced by protein folding stress are depleted of many ER proteins, which complicates their analysis.

**Figure 1.**
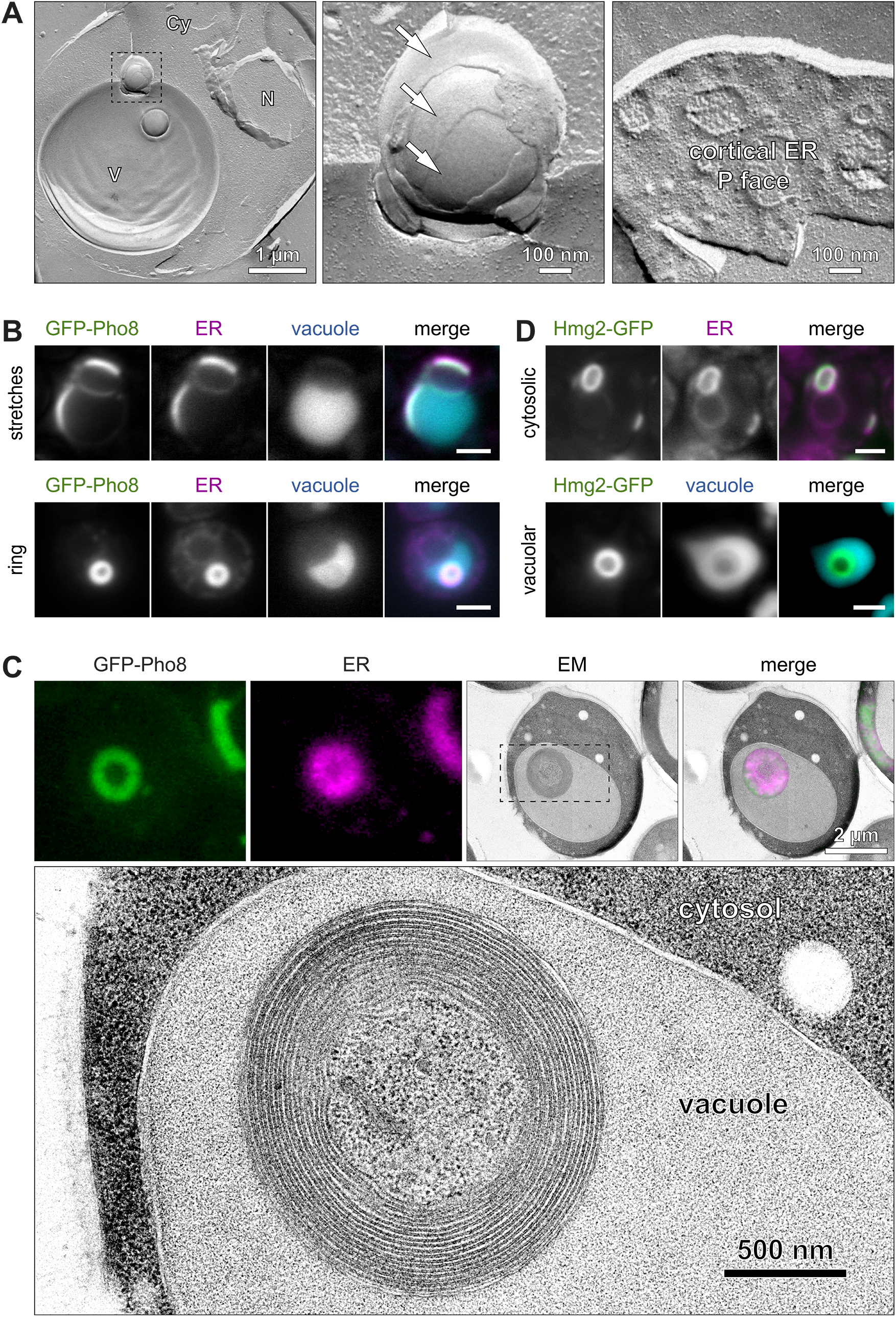
Expression of ER-resident transmembrane proteins can induce autophagy of ER whorls. **(A)** Freeze-fracture electron micrograph of wild-type yeast treated with DTT for 3 h. The left panel shows two spherical structures, one in a vacuole membrane invagination and one inside the vacuole. The middle panel shows the boxed area of the left panel at higher magnification. Arrows indicate membrane layers that are free of intramembrane particles. The right panel shows the protoplasmic fracture face (P face) of cortical ER with a high density of intramembrane particles. Cy, cytosol; N, nucleus; V, vacuole. **(B)** Fluorescence images of cells expressing GFP-Pho8 and the general ER marker Sec63-mCherry and stained with CMAC to label the vacuole. The upper and lower panel show vacuole-associated and perinuclear GFP-Pho8 stretches and an intravacuolar GFP-Pho8 ring, respectively. Scale bars: 2 *µ*m. **(C)** Correlative light and electron microscopy of cells expressing GFP-Pho8 and Sec63-mCherry. The lower panel shows the boxed area in the electron micrograph at higher magnification. The ER ring corresponds to a vacuolar whorl and the ER stretch to a perinuclear stack. **(D)** Fluorescence images of cells expressing Hmg2-GFP and either additionally expressing Sec63-mCherry or stained with CMAC. The upper and lower panel show cytosolic and vacuolar Hmg2-GFP rings, respectively. Scale bars: 2 *µ*m.

ER whorls can also be induced by high-level expression of transmembrane proteins that are genuine ER proteins or are retained in the ER because of their propensity to oligomerize (Gong et al., 1996; Koning et al., 1996; Snapp et al., 2003; Lingwood et al., 2009). We fortuitously found that the vacuolar transmembrane phosphatase Pho8, which dimerizes via its lumenal domain (Dancourt and Barlowe, 2009), became such a protein when its cytosolic N-terminus was fused to regular GFP. At high levels, GFP-Pho8 co-localized with Sec63-mCherry, indicating that it was retained in the ER, and induced brightly labelled stretches and rings (Figure 1B). No such structures formed in cells that expressed Pho8 fused to strictly monomeric GFP(L221K). Many GFP-Pho8 stretches were associated with the vacuole membrane and most GFP-Pho8 rings were inside the vacuole. Correlative light and electron microscopy demonstrated that stretches corresponded to stacked cisternal ER and that rings corresponded to whorls (Figure 1C). These whorls consisted of several layers of ribosome-free ER membrane and typically had a diameter of 1.5 *µ*m. Thus, GFP-Pho8 whorls are multilamellar, spherical and targeted by autophagy, similar to whorls formed during ER stress. However, GFP-Pho8 whorls contained various ER proteins (Figures 1B and S1C), indicating that the mechanisms that restrain protein entry into ER stress-induced whorls do not apply to them. To test whether autophagy of ER whorls induced by GFP-Pho8 reflected a common phenomenon, we examined the ER transmembrane protein Hmg2-GFP. Consistent with earlier studies, high-level expression of Hmg2-GFP yielded ER stretches at the nuclear envelope and the cell cortex, which represent ER stacks, as well as ring-shaped ER structures, which represent ER whorls (Wright et al., 1988; Koning et al., 1996; Federovitch et al., 2008). Most whorls were in the cytosol, but a small fraction were inside the vacuole (Figure 1D). Thus, ER whorls generated by expression of different ER-resident transmembrane proteins can become subject to autophagy.

To define the membrane dynamics of autophagy of ER whorls, we applied time-lapse imaging. Cells expressing GFP-Pho8 showed ER stacks that were associated with the vacuole membrane, bent towards the vacuole lumen and were converted into vacuolar whorls (Figure 2A). The time for conversion of bent stacks into vacuolar whorls was six minutes on average but showed a broad, bimodal distribution (Figure S2A). Whorls inside the vacuole were enclosed by vacuole membrane, showing that they had been taken up by microautophagy. Accordingly, electron micrographs revealed bent ER stacks in large vacuole membrane invaginations (Figure 2B). Occasionally, whorls inside the vacuole appeared as incomplete rings, suggesting that they can retain an opening (Figure S2B). Time-lapse imaging established that the microautophagic process was directional but reversible up to a late stage. Bent stacks in clearly discernible vacuole membrane invaginations, termed uptake intermediates, had a 90% chance of becoming a vacuolar whorl (Figure 2C). Yet, about 10% of uptake intermediates reverted to vacuole-associated stacks. Even structures that seemed to be vacuolar whorls could revert to uptake intermediates and then to vacuole-associated stacks or cytosolic whorls (Figure 2D). Due to limited temporal and spatial resolution, we were unable to determine whether such whorls had completed the uptake process, including vacuole membrane scission, or had simply remained in long-lived vacuole membrane invaginations. Finally, cytosolic whorls sometimes formed from ER stacks at the cell cortex (Figure S2C). Autophagy of these whorls was not observed, although we cannot rule out that it occurred with low frequency. Taken together, the use of GFP-Pho8 confirmed that autophagy of ER whorls occurs by microautophagy, revealed that stacks are precursors of whorls and showed that whorls can form during uptake into the vacuole.

**Figure 2.**
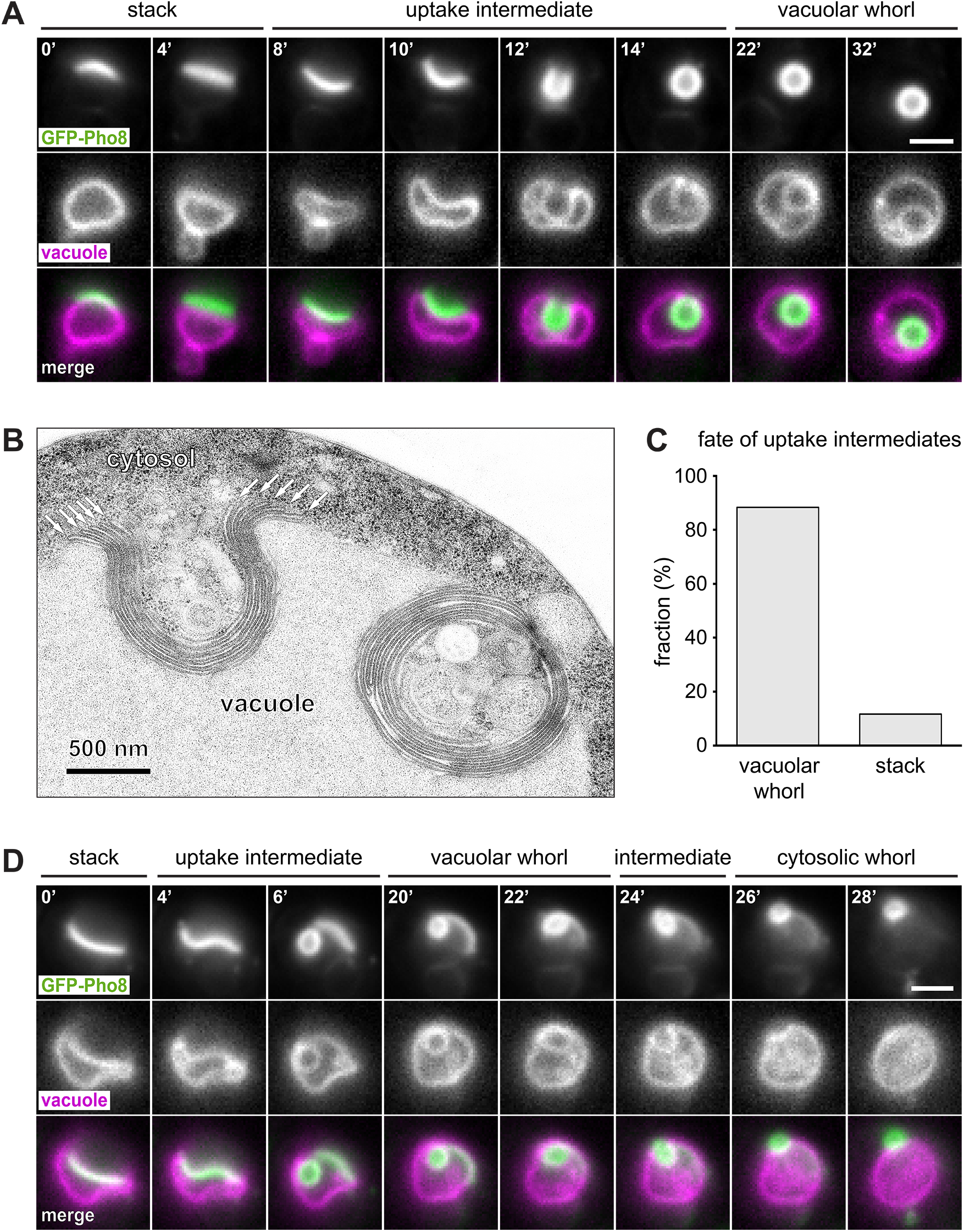
Autophagy of ER whorls occurs by microautophagy. **(A)** Individual frames from time-lapse imaging of a cell expressing GFP-Pho8 and stained with the vacuole membrane dye FM4-64. Numbers indicate the time in minutes after the start of the image sequence. Scale bar: 2 *µ*m. **(B)** Electron micrograph of a cell expressing GFP-Pho8. Arrows indicate the edges of ER cisternae that are part of a bent stack in a vacuole membrane invagination. **(C)** Fraction of uptake intermediates that were converted into vacuolar whorls or stacks. n = 43. **(D)** As in panel A. The image sequence shows formation of a vacuolar whorl that becomes a cytosolic whorl.

### Nem1-Spo7 and ESCRT proteins are key components for micro-ER-phagy

We previously discovered that autophagy of ER whorls during protein folding stress does not depend on the core autophagy machinery (Schuck et al., 2014). Likewise, autophagy of GFP-Pho8 whorls did not require the autophagy proteins Atg1, 6, 7, 8, 14 or 16 (Figures 3A and S3A). The macro-ER-phagy receptors Atg39 and 40, which link ER subdomains to the core autophagy machinery (Mochida et al., 2015), were also dispensable (Figure S3B). To identify molecular machinery for micro-ER-phagy, we conducted a genetic screen. We introduced GFP-Pho8 and Sec63-mCherry into the yeast knockout collection, visually screened for mutants with aberrant whorl formation and obtained 119 hits (Table S1). The strongest hits included the ER-resident protein phosphatase Nem1, the vacuolar phosphatidylinositol-3-phosphate 5-kinase Fab1 and several ESCRT proteins (Jin et al., 2016; Schöneberg et al., 2016; Carman and Han, 2019). Mutants lacking these proteins still generated ER stacks, but whorl formation was strongly reduced or abolished (Figures 3B and S3C).

**Figure 3.**
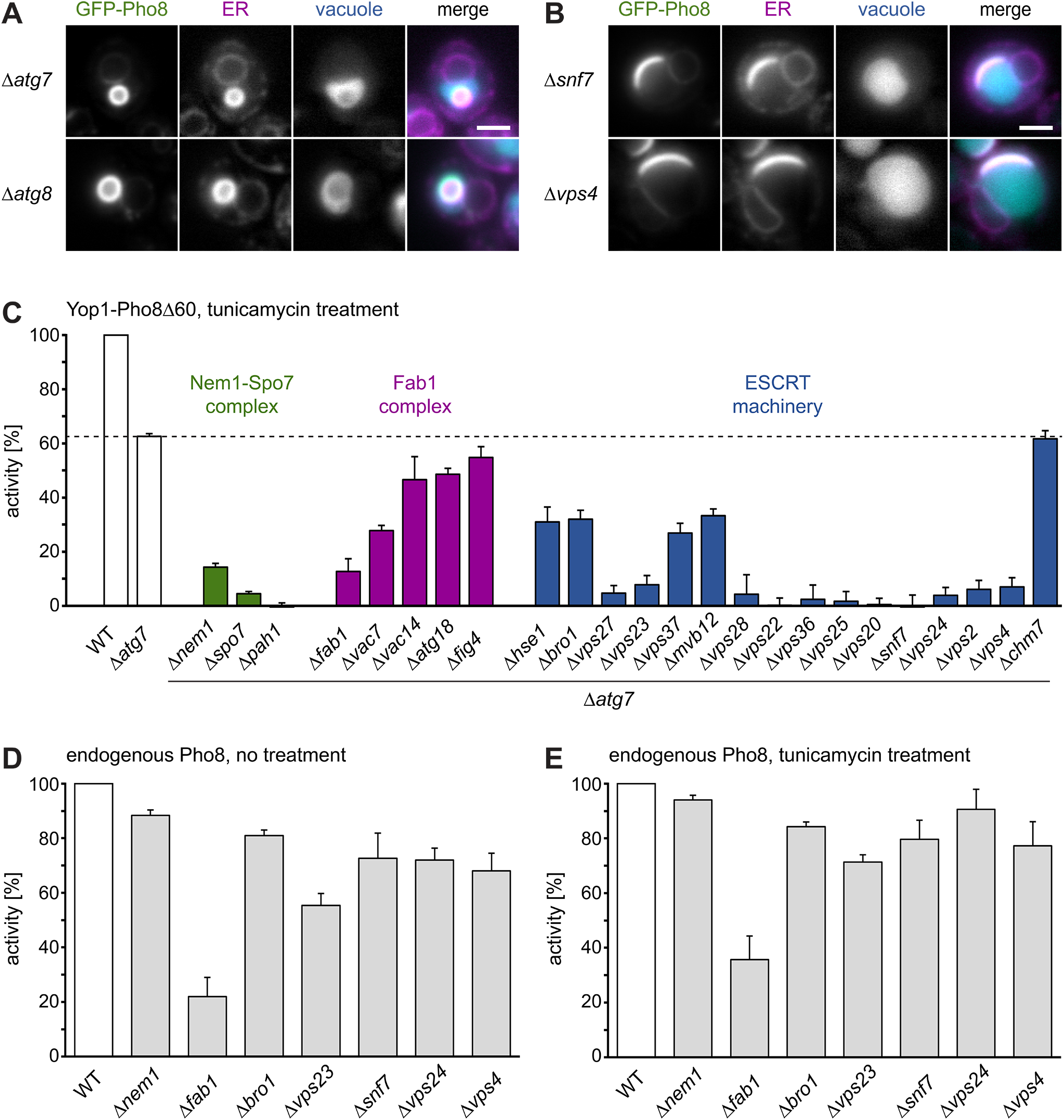
The Nem1-Spo7 complex and ESCRT proteins are key components for micro-ER-phagy. **(A)** Fluorescent images of *Δatg7* and *Δatg8* cells expressing GFP-Pho8 and the general ER marker Sec63-mCherry and stained with CMAC to label the vacuole. Scale bar: 2 *µ*m. **(B)** As in panel A but of *Δsnf7* and *Δvps4* cells. **(C)** Relative activity of the ER-phagy reporter Yop1-Pho8Δ60 in tunicamycin-treated cells. Background activity determined in *Δpep4* mutants was subtracted from the activity in all other strains. The dotted line indicates the activity in *Δatg7* cells. Mean ± SEM, n ≥ 3. **(D)** Relative activity of endogenous Pho8 in untreated cells. Background activity determined in *Δpho8* mutants was subtracted from the activity in all other strains. Mean ± SEM, n = 3. **(E)** As in panel D but after tunicamycin treatment.

To test whether these mutants were also defective in ER-phagy during protein folding stress, we employed a previously established biochemical assay. This assay is based on the phosphatase domain of Pho8, called Pho8Δ60, fused to the ER membrane protein Yop1 as part of the ER-phagy reporter Yop1-Pho8Δ60 (Schuck et al., 2014). In contrast to endogenous Pho8, in which the phosphatase domain is lumenal, fusion to Yop1 tethers Pho8Δ60 to the cytosolic surface of the ER (Figure S4A). Cytosolic Pho8Δ60 has low catalytic activity due to an autoinhibitory C-terminal propeptide. This propeptide is cleaved off only upon autophagic transport into the vacuole, yielding an active enzyme. Cells were treated with tunicamycin to induce ER-phagy and Yop1-Pho8Δ60 activity was measured. On a technical note, DTT disrupts vacuolar proteolysis, blocks activation of Yop1-Pho8Δ60 and therefore cannot be employed in this assay (Schuck et al., 2014). *Δpep4* mutants, in which vacuolar proteolysis is disrupted genetically, were used to determine background phosphatase activity. *Δatg7* mutants, in which the core autophagy machinery and hence macroautophagy is disabled, retained 60% of wild-type activity, consistent with our earlier data (Figure 3C; Schuck et al., 2014). Candidate genes were tested in a *Δatg7* background to exclude contributions from macro-ER-phagy. Additional removal of Nem1, its binding partner Spo7 or its substrate Pah1 strongly reduced ER-phagy. The same was true for Fab1 and its activator Vac7, whereas elimination of the Fab1 regulators Vac14, Atg18 or Fig4 had little effect. Removal of ESCRT-0, - I, -II, -III proteins, Bro1 or Vps4 reduced or blocked ER-phagy. Removal of Chm7, an ESCRT protein important for nuclear envelope quality control (Webster et al., 2016), had no effect. Hence, genes needed for microautophagy of GFP-Pho8 whorls are also required for micro-ER-phagy induced by protein folding stress.

To address the possibility that the observed ER-phagy defects result from general vacuole dysfunction and impaired removal of the autoinhibitory propeptide from Yop1-Pho8Δ60, we measured the activity of endogenous Pho8. Endogenous Pho8 is activated by the same proteolytic processing as Yop1-Pho8Δ60 and therefore is an ideal control. *Δfab1* mutants showed strongly reduced Pho8 activity at steady state and after tunicamycin treatment (Figures 3D and 3E), suggesting that indirect effects contribute to their ER-phagy phenotype. In contrast, *Δnem1* and ESCRT mutants showed almost normal Pho8 activity at steady-state and after tunicamycin treatment. Therefore, and in contrast to what we suspected earlier (Schuck et al., 2014), vacuole functions relevant to Yop1-Pho8Δ60 activation are largely intact in *Δnem1* and ESCRT mutants, indicating that Yop1-Pho8Δ60 can faithfully report on their ER-phagy activity. These results confirm that the Nem1-Spo7 complex and ESCRT proteins are genuine components of the micro-ER-phagy machinery.

### Macro- and micro-ER-phagy have distinct molecular requirements

Next, we aimed to delineate the contributions of the core autophagy machinery, the Nem1-Spo7 complex and ESCRT proteins to different autophagy pathways, namely to non-selective macroautophagy, macro-ER-phagy and micro-ER-phagy.

First, we measured non-selective autophagy upon nitrogen starvation using the soluble cytosolic reporter cyto-Pho8Δ60 (Figure S4A; Noda et al., 1995). Under these conditions, autophagy of cytosol occurs, primarily or exclusively, by macroautophagy (Baba et al., 1994). Cyto-Pho8Δ60 activity was abolished by deletion of *ATG7*, consistent with a strict requirement for the core autophagy machinery (Figure 4A; Mizushima et al., 1998). Deletion of *NEM1* or *SPO7* caused only minor defects, but deletion of the ESCRT components *VPS23*, *SNF7* or *VPS4* reduced non-selective macroautophagy by >50%. These defects in ESCRT mutants could be due to a role of ESCRT proteins in autophagosome closure. Alternatively, they could reflect that ESCRT proteins are needed to maintain vacuole function during starvation (Müller et al., 2015). Accordingly, endogenous Pho8 activity during nitrogen starvation was reduced by half in ESCRT mutants (Figure S4B).

**Figure 4.**
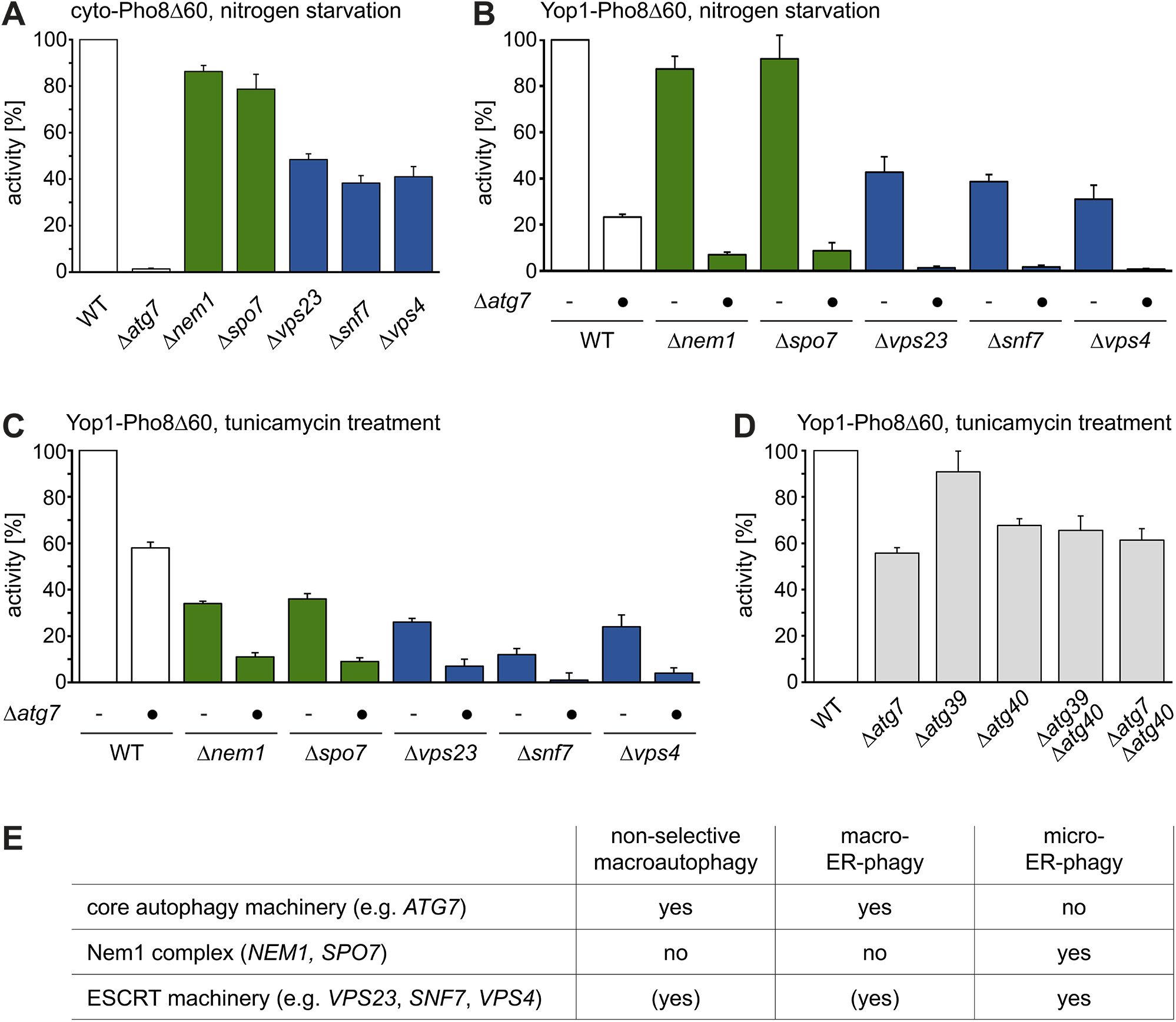
Macro- and micro-ER-phagy are parallel pathways with distinct molecular requirements. **(A)** Relative activity of cyto-Pho8Δ60 after nitrogen starvation, which reports on non-selective macroautophagy of cytosol. Mean ± SEM, n ≥ 3. **(B)** Relative activity of the ER-phagy reporter Yop1-Pho8Δ60 after nitrogen starvation. Mean ± SEM, n ≥ 3. **(C, D)** As in panel B but after tunicamycin treatment. **(E)** Summary of the requirements of different autophagy pathways. ‘(yes)’ indicates that a component is needed for maximal efficiency of an autophagy pathway.

Second, we measured ER-phagy upon nitrogen starvation using Yop1-Pho8Δ60. Deletion of *ATG7* reduced starvation-induced ER-phagy by 75% (Figure 4B). Since macroautophagy is absent in *Δatg7* cells (Mizushima et al., 1998), a different mechanism must account for at least 25% of ER turnover under these conditions. Single deletion of *NEM1* or *SPO7* mildely reduced starvation-induced ER-phagy, and deletion of *ATG7* together with either *NEM1* or *SPO7* caused an almost complete block. Hence, disruption of the core autophagy machinery and the Nem1-Spo7 complex additively inhibited starvation-induced ER-phagy. This result suggests that the two modules act in separate pathways, which likely correspond to macro- and micro-ER-phagy. Single deletion of ESCRT components reduced starvation-induced ER-phagy by >60%, and combined disruption of the core autophagy machinery and ESCRT components abolished it. As shown above Atg7-independent micro-ER-phagy is blocked in ESCRT mutants but contributes only 25% of ER turnover during nitrogen starvation. Therefore, the magnitude of the defects in ESCRT single mutants shows that ESCRT proteins are also needed for efficient macro-ER-phagy.

Third, we analyzed ER stress-induced ER-phagy using Yop1-Pho8Δ60. As seen before, deletion of *ATG7* reduced ER-phagy upon tunicamycin treatment by 40% (Figures 3C and 4C). Thus, Atg7-independent micro-ER-phagy accounts for 60% of ER turnover under these conditions. Single deletion of *NEM1* or *SPO7* eliminated roughly 60% of ER-phagy activity and combined deletion of *ATG7* and either *NEM1* or *SPO7* reduced ER-phagy by about 90%. A parsimonious explanation for these results is that the core autophagy machinery and the Nem1-Spo7 complex function largely in parallel also during ER stress. Single deletion of ESCRT components strongly reduced ER-phagy, which can again only be explained by impairment of both Atg7-independent micro-ER-phagy and Atg7-dependent macro-ER-phagy.

Last, we tested Atg39 and Atg40, which mediate macroautophagy of perinuclear and peripheral ER, respectively (Mochida et al., 2015). Deletion of *ATG40* reduced Yop1-Pho8Δ60 activity (Figures 4D and S4C), consistent with localization of Yop1 to the peripheral ER (Voeltz et al., 2006). However, *ATG40* deletion had no effect in *Δatg7* cells. Hence, Atg40 is dispensable for Atg7-independent micro-ER-phagy.

Collectively, these results indicate that macro- and micro-ER-phagy are parallel pathways. Their relative contributions to ER turnover depend on how ER-phagy is triggered and their molecular requirements are overlapping yet distinct. The core autophagy machinery is essential for macro-ER-phagy but dispensable for micro-ER-phagy. The Nem1-Spo7 complex is primarily involved in micro-ER-phagy, although it makes a minor contribution to macro-ER-phagy. ESCRT proteins are essential for micro-ER-phagy and are additionally needed for efficient macro-ER-phagy and non-selective macroautophagy (Figure 4E).

### ESCRT proteins mediate vacuole membrane dynamics during micro-ER-phagy

To understand the role of ESCRT proteins in micro-ER-phagy, we performed time-lapse imaging of wild-type, *Δvps23*, *Δsnf7* and *Δvps4* cells. GFP-Pho8 expression levels and the fraction of cells with GFP-Pho8 structures were similar in wild-type and ESCRT mutant cells (Figures S5A and 5A). GFP-Pho8 structures were classified as stacks (at the vacuolar membrane, nuclear envelope or cell periphery), cytosolic whorls, uptake intermediates (bent stacks in vacuole membrane invaginations) or vacuolar whorls (circular and inside the vacuole). In wild-type cells, 80% of GFP-Pho8 structures were classified as stacks, <5% as cytosolic whorls, <5% as uptake intermediates and 15% as vacuolar whorls (Figure 5B). The fraction of cytosolic whorls was increased rather than decreased in *Δvps23*, *Δsnf7* and *Δvps4* cells, showing that ESCRT proteins are not required for whorl formation in the cytosol. In contrast, the fraction of vacuolar whorls was sharply reduced to 2% or less, with *Δsnf7* cells showing the most severe phenotype. Rare instances of vacuolar whorl formation in ESCRT mutants showed that the process occurred via the same morphological stages as in wild-type cells (Figure S5B). Since stacks at the vacuole membrane are precursors for vacuolar whorls, we asked whether the association of ER stacks with the vacuole was affected in ESCRT mutants. The fraction of vacuole-associated stacks was 70% in wild-type cells but only 30% in cells lacking Snf7 (Figure 5C). Overall, these results show that ESCRT proteins are major mediators of the microautophagic uptake of ER into the vacuole.

**Figure 5.**
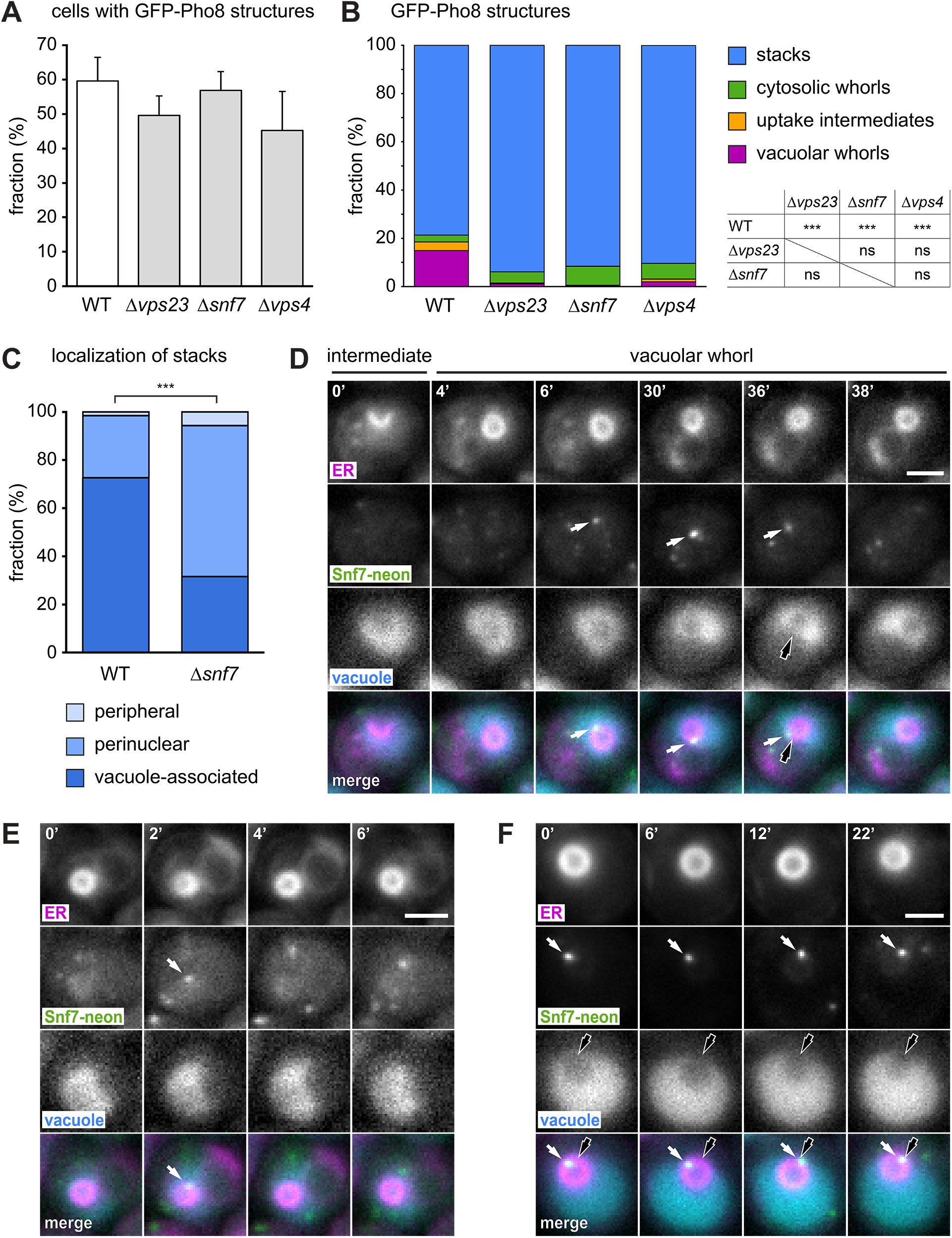
ESCRT machinery mediates vacuole membrane dynamics during micro-ER-phagy of GFP-Pho8 structures. **(A)** Fraction of wild-type (WT), *Δvps23*, *Δsnf7* and *Δvps4* cells with GFP-Pho8 structures. Mean ± SEM, n ≥ 3. **(B)** Fraction of GFP-Pho8 structures classified as stacks, cytosolic whorls, uptake intermediates or vacuolar whorls. The number of GFP-Pho8 structures in WT, *Δvps23*, *Δsnf7* and *Δvps4* cells was 562, 345, 367 and 449, respectively. The table shows the results of Chi-Square tests for independence for all pairs of strains. ***, highly significant (P < 0.00001); ns, not significant (P > 0.05). **(C)** Fraction of stacks classified as peripheral, perinuclear or vacuole-associated. The number of stacks in WT and *Δsnf7* cells was 252 and 351, respectively. The observed distributions in the two strains were analyzed with a Chi-Square test for independence. ***, highly significant (P < 0.00001). **(D)** Fluorescence images of a cell expressing non-fluorescent GFP(Y66F)-Pho8, Snf7-mNeonGreen and the ER marker mCherry-Ubc6, and stained with the vacuole dye CMAC. Numbers indicate the time in minutes after the start of the image sequence. White arrows indicate a Snf7-mNeonGreen punctum adjacent to an ER whorl. Black arrow point to a vacuole membrane invagination. Scale bar: 2 *µ*m. **(E)** As in panel D, illustrating transient appearance of Snf7 at an ER whorl. **(F)** As in panel D, illustrating orientation of a Snf7 punctum towards the neck of a vacuole membrane invagination. In this series, mNeonGreen fluorescence was adjusted differently for different time points to compensate signal decay over time.

To define at which steps during vacuolar uptake of ER the ESCRT machinery acts, we imaged Snf7, the major component of ESCRT-III assemblies (Schöneberg et al., 2016; McCullough et al., 2018). Tagged Snf7 is non-functional but can serve as a valid tracer of ESCRT action when present alongside endogenous Snf7 (Adell et al., 2017). To minimize the risk of dominant negative effects, we introduced, into wild-type cells, an expression construct for Snf7-mNeonGreen under the *VPS24* promoter, which is about three-fold weaker than the *SNF7* promoter (Figure S5C). In addition, we made GFP-Pho8 compatible with visualization of Snf7-mNeonGreen by mutating its chromophore, yielding non-fluorescent GFP(Y66F)-Pho8. Finally, we employed mCherry-Ubc6 as an ER marker, stained the vacuole with CMAC, triggered micro-ER-phagy by expression of GFP(Y66F)-Pho8 and performed time-lapse imaging. Snf7 puncta abruptly appeared adjacent to late ER uptake intermediates, remained associated for some time, and rapidly disappeared, leaving behind vacuolar whorls free of Snf7 (Figure 5D, white arrows, and Movie 1). The duration of Snf7 appearance at whorls was variable and occasionally as short as a single movie frame (Figure 5E). Inspection of optical sections covering entire GFP-Pho8 structures confirmed that Snf7 puncta associated with the periphery of ER whorls and not their interior (Movie 1). Furthermore, Snf7 puncta juxtaposed to ER whorls were typically oriented towards the neck of vacuole membrane invaginations (Figure 5D, black arrow, Figure 5F and Movie 2). This transient association of Snf7 with sites of ER uptake suggests that the ESCRT machinery is directly involved in a late step of uptake but is released before uptake is complete. Snf7 puncta appeared close to ER stacks after they had already been bent into whorls. This observation indicates that Snf7 acts subsequent to vacuole membrane bending and ER whorl formation, and at the time when scission of the vacuole membrane occurs.

The above results suggested a new interpretation for data we had reported earlier. Specifically, we had previously found vacuolar whorls in electron micrographs of DTT-treated *Δvps23* and *Δvps4* cells and concluded that the ESCRT machinery is not essential for micro-ER-phagy (Schuck et al., 2014). Given the role of ESCRT proteins in vacuole membrane scission, we tested the possibility that whorls induced by ER stress are trapped in vacuole membrane invaginations in ESCRT mutants.

Consistent with our previous findings, we observed vacuolar whorls in electron micrographs of DTT-treated *Δsnf7* cells, which were indistinguishable from vacuolar whorls in wild-type cells (Figure 6A). However, analysis of serial thin sections revealed that 10 out of 10 vacuolar whorls in *Δsnf7* cells were still connected to the cytosol and had not completed the uptake process (Figures 6B and 6C). In contrast, 8 out of 15 whorls in wild-type cells were completely disconnected from the cytosol (Figures 6B and 6D). It is improbable that this difference in the proportions of connected and disconnected whorls in wild-type and *Δsnf7* cells arose by chance (p = 0.008 according to Fisher’s exact test, Figure 6B). Instead, it likely reflects slowed or blocked microautophagic uptake of ER whorls in *Δsnf7* cells. These observations indicate that ESCRT proteins are needed for completion of micro-ER-phagy also during ER stress.

**Figure 6.**
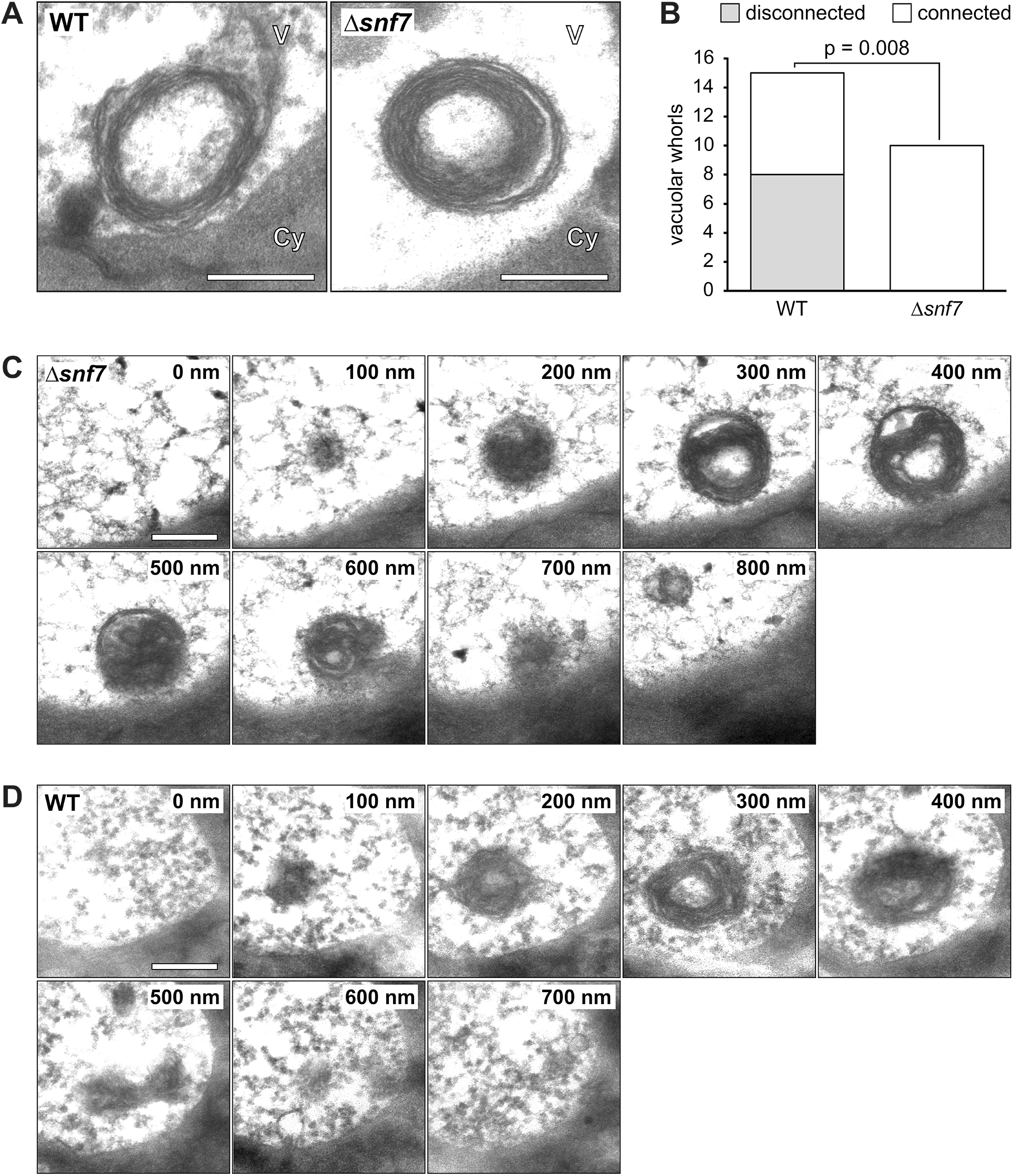
ESCRT machinery is required for completion of microautophagy of ER whorls during ER stress. **(A)** Electron micrographs of wild-type and *Δsnf7* cells treated with DTT for 3 h. Cy, cytosol; V, vacuole. Scale bars: 300 nm. **(B)** Quantification of whorls that are disconnected from the cytosol or still connected to the cytosol. The observed distributions in the two strains were analyzed with a Fisher’s exact test. **(C)** Electron micrographs of serial thin sections of a *Δsnf7* cell treated with DTT for 3 h. Note that the whorl appears disconnected from the cytosol in some sections (e.g. section 300 nm) but still is connected to the cytosol (see sections 600 nm and 700 nm). Scale bar: 300 nm. **(D)** As in panel C but of a wild-type cell. The vacuolar whorl is completely disconnected from the cytosol.

## DISCUSSION

We have shown that micro-ER-phagy in yeast involves the conversion of stacked cisternal ER into whorls during uptake into the vacuole. In contrast to macro-ER-phagy, micro-ER-phagy does not need the core autophagy machinery. Instead, the process requires the Nem1-Spo7 complex and ESCRT machinery, which mediates vacuole membrane scission during microautophagic uptake of ER (Figure 7).

**Figure 7.**
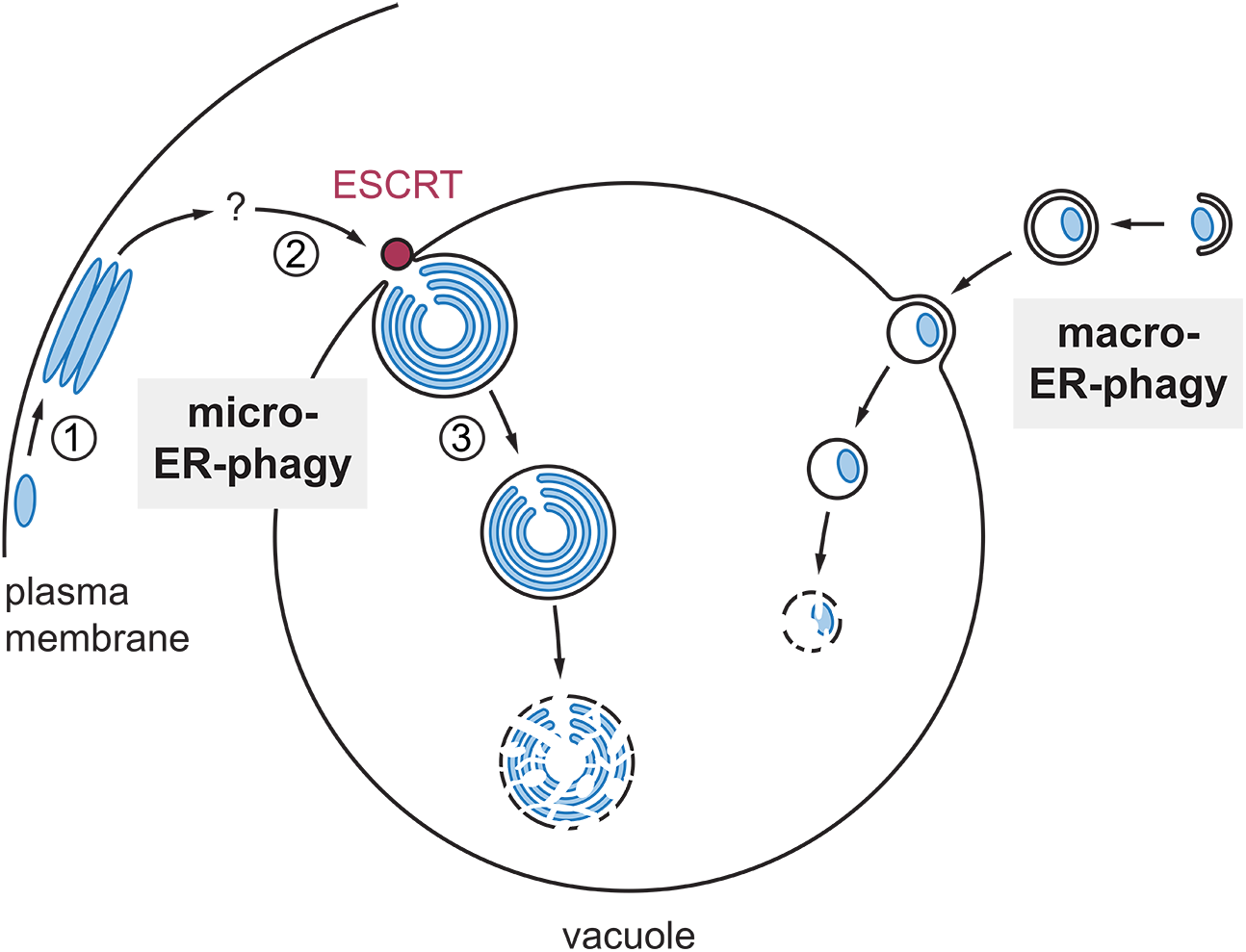
Model of ER-phagy. During micro-ER-phagy, portions of the ER are reorganized to form stacks (1). ER stacks are bent, recognized by the vacuole, taken up into vacuole membrane invaginations, converted into whorls and detached from the remainder of the ER (2). The order of these events is uncertain, as indicated by the question mark. ESCRT proteins are recruited to sites of ER whorl uptake and mediate scission of the vacuole membrane (3). In parallel, macro-ER-phagy can also selectively target the ER, which is mediated by the core autophagy machinery.

In this study, we employed a fusion protein, GFP-Pho8, to trigger microautophagy of ER whorls. Compared with whorls induced by ER stress, GFP-Pho8 whorls have an approximately three-fold larger diameter (Schuck et al., 2014) and do not recapitulate the intriguing depletion of many ER proteins. Nevertheless, the ER protein content of GFP-Pho8 whorls, their size, slow kinetics of vacuolar uptake and stability inside the vacuole made them a powerful tool for investigating micro-ER-phagy. Importantly, the genes required for microautophagy of GFP-Pho8 whorls are also needed for micro-ER-phagy induced by ER stress, as observed by electron microscopy and measured by the Yop1-Pho8Δ60 reporter. Considering the depletion of proteins from stress-induced whorls, it seems paradoxical that the Yop1-Pho8Δ60 reporter is useful at all. Moreover, Yop1 preferentially localizes to high-curvature tubular ER (Voeltz et al., 2006), whereas whorls consist of low-curvature ER cisternae. However, the presence of an opening in GFP-Pho8 whorls indicates that the edges of the bent ER sheets that make up ER whorls remain unfused. Hence, GFP-Pho8 whorls, and presumably also stress-induced whorls, contain high-curvature sheet edges at their opening, which makes Yop1-Pho8Δ60 a suitable biochemical reporter for autophagy of otherwise protein-depleted ER whorls.

ESCRT proteins mediate membrane remodelling and scission in various processes, for example during biogenesis of 25-nm intralumenal vesicles at late endosomes in yeast (Henne et al., 2013; Adell et al., 2017). However, ESCRT proteins can also participate in the scission of 1-µm wide membrane buds, for instance during cytokinesis, and can function at the vacuole membrane in yeast (Henne et al., 2013; Zhu et al., 2017). The ESCRT machinery appears to be remarkably flexible and it remains to be uncovered how it is adapted for micro-ER-phagy. Importantly, we provide evidence that ESCRT-III is directly involved in vacuole membrane scission. The impaired vacuole association of GFP-Pho8 stacks and the reduced formation of uptake intermediates in ESCRT mutants could reflect indirect effects or roles of ESCRT proteins also in these earlier steps. Residual formation of bent stacks and whorls occurs in ESCRT mutants, pointing to additional mechanisms. Intriguingly, the Nem1-Spo7 complex appears to be generally involved in microautophagy (Tsuji et al., 2017; Rahman et al., 2018) and is required for formation of vacuole membrane domains (Toulmay et al, 2013). Such domains are sites of microautophagy of lipid droplets (Wang et al., 2014; Tsuji et al., 2017) and the line tension at domain boundaries may drive membrane bending (Lipowsky, 2002). Whether domain-induced budding is important for micro-ER-phagy remains to be elucidated. Furthermore, identifying the determinants that enable recognition of ER stacks and their segregation from regular ER for autophagic degradation is a key challenge.

The extensive reorganization of membranes during the formation of ER whorls raises many interesting questions. Exclusion of certain ER proteins from stacks and whorls has been reported before (Gong et al., 1996; Koning et al., 1996; Okiyoneda et al., 2004; Federovitch et al., 2008), but our freeze-fracture electron microscopy data are the first to indicate that ER whorls can be almost free of transmembrane proteins. The underlying mechanisms are unknown but may serve to spare well-folded proteins from autophagic degradation. Whorl formation has been proposed to involve zippering of ER membranes by proteins that self-interact in trans (Gong et al., 1996; Snapp et al., 2003). Such zippering could explain the formation of GFP-Pho8 whorls but it is unclear how it might occur for the protein-depleted whorls induced by ER stress. Another open issue is how stacked ER membranes become associated with the vacuole membrane. The yeast vacuole is motile and has access to the entire cytoplasm, so that association of ER and the vacuole can take place at the cell cortex (Schuck et al, 2014). Furthermore, the force that bends stacks into micrometer-sized whorls remains to be identified. As mentioned above, one possibility is that this force is provided, at least in part, by domain-induced budding of the vacuole membrane. Last, the ER taken up into the vacuole needs to be detached from the remainder of the ER, yet it is unknown how ER fission might occur.

Our study opens up micro-ER-phagy for mechanistic analysis and has general implications for microautophagy and ER homeostasis. Microautophagy has remained enigmatic, in part because it is an umbrella term for distinct pathways. However, our findings support a unifying view that ESCRT proteins have direct roles in many, perhaps all, microautophagy pathways. By interfering with the metazoan homologues of ESCRT proteins and the Nem1-Spo7 complex, it is now possible to explore the possible conservation of micro-ER-phagy (Omari et al., 2018) and further investigate the physiological functions of the process. Autophagy eliminates excess ER in both yeast and mammalian cells (Bohlender and Weibel, 1973; Schuck et al., 2014; Fumagalli et al., 2016), and the finding that proteins can be segregated from ER subdomains that give rise to whorls emphasizes the remarkable selectivity of micro-ER-phagy. In addition, the identification of molecular machinery for micro-ER-phagy will allow to determine why macro- and micro-ER-phagy coexist and are differentially employed depending on conditions. Finally, the medical importance of ER-phagy in viral infection, neuropathy and cancer is just beginning to emerge (Grumati et al., 2018), and understanding both macro- and micro-ER-phagy will help uncover how autophagy promotes organellar and hence organismal health.

## ACKNOWLEDGEMENTS

We thank Randy Hampton, Benoit Kornmann and David Teis for plasmids, the ZMBH Imaging Facility, the Flow Cytometry and FACS facility at the ZMBH and the Heidelberg University Electron Microscopy Core Facility for support, Georg Borner and David Teis for discussion and advice, and Frauke Melchior, Anne Schlaitz and all members of the Schuck lab for comments on the manuscript. This work was funded by a PhD fellowship from the Heidelberg Biosciences International Graduate School to J.A.S. and grant EXC 81 from the Deutsche Forschungsgemeinschaft.

## AUTHOR CONTRIBUTIONS

Conceptualization, J.A.S. and S.S.; Software, P.W.B. and J.P.S.; Investigation, C.F., D.P., G.R., J.A.S., K.S., J.P.S., S.S. and T.T.; Resources, M.K.; Writing – Original Draft, J.A.S. and S.S.; Writing – Review & Editing, all authors.

## MATERIALS AND METHODS

### Plasmids

Plasmids used in this study are listed in Table S2 and sequences of oligonucleotides for plasmid construction are given in Table S3. To generate pRS406-P_GPD_-GFP-Pho8, pRS406-P_GPD_-mCherry-Ubc6 was amplified with primers p406 rev GPD overhang/CYCterm fw Pho8 overhang and ligated, by means of the NEBuilder HiFi DNA assembly master mix (New England Biolabs, Ipswitch, Massachusetts), with P_GPD_-GFP-Pho8 amplified with primers GPD-GFPPho8 fw/GFPPho8 rev from genomic DNA of yeast strain SSY1094. pRS406-P_GPD_-mGFP-Pho8 containing monomeric GFP(L221K) was generated from pRS406-P_GPD_-GFP-Pho8 by site-directed mutagenesis with primers yeGFP L221K fw/rev. To generate pRS406-P_GAL_-GFP-Pho8, pRS406-P_GPD_-GFP-Pho8 was amplified with primers p406 GFPPho8 fw/p406 GFPPho8 rev and ligated with the *GAL* promoter sequence amplified from pRS416-P_GAL_ (Mumberg et al., 1994) with primers GAL fw overhang/GAL rev overhang. To generate pRS405-P_GAL_-GFP-Pho8, pRS406-P_GAL_-GFP-Pho8 was amplified with primers p406 no URA fw/p406 no URA rev to remove the *URA3* gene and ligated with the *LEU2* gene amplified from pRS415 (Sikorski and Hieter, 1989) with primers LEU2 prom fw overhang/LEU2 cds rev overhang. To generate pRS406-P_GAL_-Hmg2(K6R)-GFP, Hmg2(K6R)-GFP was first amplified from Yip-P_GAL_-Hmg2(K6R)-GFP (Federovitch et al., 2008) with primers Xba-Hmg2/GFP-Bam and cloned into the XbaI/BamHI site of pRS415-P_ADH_ (Mumberg et al., 1995), yielding pRS415-P_ADH_-Hmg2(K6R)-GFP. The Hmg2(K6R)-GFP-T_CYC_ cassette was then excised from pRS415-P_ADH_-Hmg2(K6R)-GFP with XbaI and KpnI and cloned into the XbaI/KpnI site of pRS406-P_GAL_-GFP-Pho8, yielding pRS406-P_GAL_-Hmg2(K6R)-GFP. To generate pRS404-P_GPD_-mCherry-Ubc6, the expression cassette for mCherry fused to the C-terminal 18 amino acids of Ubc6 was excised from pRS406-P_GPD_-mCherry-Ubc6 with AatIII/NaeI and cloned into the AatIII/NaeI site of pRS404 (Sikorski and Hieter, 1989). To generate pRS416-P_SNF7_-Snf7-LAP-mNeonGreen, pRS416-Snf7-LAP-eGFP (Adell et al., 2017) was amplified with primers Snf7 term fw neon overhang/LAP rev neon overhang to remove the eGFP coding sequence and ligated with mNeonGreen coding sequence amplified from pFA6a-mNeonGreen-kanMX6 with primers neon fw/rev. To generate pRS416-P_VPS24_-Snf7-LAP-mNeonGreen, pRS416-P_SNF7_-Snf7-LAP-mNeonGreen was amplified with primers Snf7 VPS24pr overhang fw/backbone VPS24pr overhang rev and ligated with the *VPS24* promoter amplified from genomic DNA of strain SSY122 with primers VPS24pr fw/rev. pRS405-P_GAL_-GFP(Y66F)-Pho8 was generated from pRS405-P_GAL_-GFP-Pho8 by site-directed mutagenesis with primers GFP(Y66F) fw/rev.

### Yeast strain generation

Strains were derived from W303 MATa unless indicated otherwise and are listed in Table S4. Gene tagging and deletion was done with PCR products (Longtine et al., 1998; Janke et al., 2004). After inserting plasmids into the genome, clones with the desired number of integrations were selected by flow cytometry. To generate SSY1653, *SEC63* in strain Y7092 (Tong and Boone, 2007) was tagged with mCherry, yielding SSY1646. Next, to enable genomic integration of multiple copies of the P_GAL1_-GFP-Pho8 expression cassette, the deleted *URA3* coding sequence in SSY1646 was restored by integration of the wild-type *URA3* gene amplified from pRS406-P_GAL1_-GFP-Pho8 with primers URA3pr fw overhang/URA3ter rev overhang. The newly introduced wild-type *URA3* coding sequence was then replaced with the *ura3-1* allele amplified from SSY122 genomic DNA with primers URA3 cds fw and URA3 cds rev. Counterselection with 5-fluoroorotic acid yielded strain SSY1648. Finally, cells were transformed with XcmI-linearized pRS406-P_GAL1_-GFP-Pho8 and a clone that had integrated four copies of the plasmid was identified by flow cytometry.

### Growth conditions

Strains were cultured at 30°C in YPD or SC medium. YPD consisted of 1% yeast extract (Becton Dickinson, Heidelberg, Germany), 2% peptone (Becton Dickinson) and 2% glucose (Merck, Darmstadt, Germany). SC consisted of 0.7% yeast nitrogen base (Sigma, Taufkirchen, Germany), amino acids and either 2% glucose, 2% raffinose (Sigma) or 1% raffinose/2% galactose (AppliChem, Darmstadt, Germany). Raffinose and galactose were sterile-filtered rather than autoclaved to avoid hydrolysis or caramelization. SC media containing glucose, raffinose or raffinose/galactose are referred to as SCD, SC-raf or SC-raf/gal, respectively.

### Electron microscopy

For conventional chemical fixation, cells were grown to OD_600_ = 0.5 in YPD medium, and either diluted to OD_600_ = 0.1 and left untreated, or diluted to OD_600_ = 0.2 and treated with 8 mM DTT (Roche, Mannheim, Germany). Cells were grown for another 3 h, ten ODs of cells were harvested by centrifugation at 1,000 xg at room temperature for 5 min, resuspended in 1 ml fixative (40 mM KHPO_4_ pH 7.0 containing 1% EM grade glutaraldehyde and 0.2% EM grade formaldehyde from Electron Microscopy Sciences (EMS, Munich, Germany) and incubated for 3 min.

Cells were pelleted at 1,500 xg for 2 min, resuspended in 1 ml fresh fixative and incubated on ice for 50 min. At room temperature, cells were washed for 3 × 5 min with water, resuspended in freshly prepared 2% KMnO_4_ (EMS) and incubated for 3 min. Cells were pelleted at 1,500 xg for 2 min, resuspended in 1 ml fresh 2% KMnO_4_ and incubated for 45 min. Cells were dehydrated by consecutive incubations for 5 min each in 50%, 70%, 80%, 90%, 95% and 100% (v/v) ethanol (EMS). After a final wash in water-free ethanol, cells were washed for 3 × 10 min with propylene oxide (EMS) and embedded in epon resin (Embed-812, EMS). Cells were infiltrated at room temperature with 25% resin in propylene oxide for 1 h, followed by 50% resin for 2 h, 75% resin for 4 h, pure resin at 30°C for 1 h, pure resin at 30°C for 2 × 12 h and pure resin with 3% accelerator (BDMA, EMS) at 30°C for 12 h. Resin was cured at 60°C for two days, 70 - 100 nm thin sections were cut and stained with 2% aqueous uranyl acetate for 5 min followed by Reynolds’ lead citrate for 3 min.

Images were acquired with a Jeol JEM-1400 80kV transmission electron microscope (Jeol, Tokyo, Japan) equipped with a TemCam F416 digital camera (TVIPS, Gauting, Germany). Serial sections of 100 nm thickness were used for quantification of vacuolar whorls that were disconnected from or still connected to the cytosol.

Randomly renamed series of electron micrographs showing vacuolar whorls in wild-type and *Δsnf7* cells were independently scored as disconnected or connected by two individuals who did not know the identity of the cells. If no consensus was reached, the series in question was discarded as inconclusive.

For freeze-fracture electron microscopy, cells were grown and treated with DTT as above. EM grids impregnated with cells were sandwiched between a 20 μm–thick copper foil and a flat aluminum disc (Engineering Office M. Wohlwend, Sennwald, Switzerland) and frozen with an HPM 010 high-pressure freezer (Leica Microsystems, Wetzlar, Germany) as described (Tsuji et al., 2017). The frozen specimens were transferred to the cold stage of a Balzers BAF 400 apparatus and fractured at −105°C under a vacuum of approximately 1 × 10^-6^ mbar. Replicas were made by electron-beam evaporation of platinum-carbon (1 – 2 nm) at an angle of 45° followed by carbon (10 – 20 nm) at an angle of 90°. The replicas were treated with household bleach to remove biological materials, picked up on formvar-coated EM grids, and observed with a Jeol JEM-1011 EM (Jeol) operated at 100 kV. Digital images were captured using a CCD camera (Gatan, Pleasanton, California).

For correlative light and electron microscopy, strain SSY1425 was grown to OD_600_ = 6.6 in SCD, diluted to OD_600_ = 0.2 in fresh medium and grown for another 1.5 h. Cells were collected by filtration, transferred into 0.2 mm deep aluminium carriers (Engineering Office M. Wohlwend, Sennwald, Switzerland) and frozen with a HPM010 high-pressure freezer (Bal-Tec, Liechtenstein). Freeze substitution was done with a EM AFS2 automated freeze substitution system (Leica Microsystems, Wetzlar, Germany). Samples were kept in dry acetone containing 0.1% uranyl acetate at −90°C for 24 h. The temperature was increased to −45°C at a rate of 5°C/h and maintained at −45°C for another 5 h. Samples were washed with dry acetone for 3 × 10 min, incubated in 10% Lowicryl resin in dry acetone for 2 h and in 25% resin for another 2 h. Samples were transferred to 50% resin, the temperature was increased to −35°C at a rate of 5°C/h and maintained at 35°C for another 2 h. Samples were transferred to 75% resin, the temperature was increased to −25°C at a rate of 5°C/h and maintained at −25°C for another 2 h. Samples were incubated in 100% resin at −25°C for 3 × 10 h. UV polymerization was done at −25°C for 24 h, the temperature was increased to 20°C under UV light within 9 h and polymerization was continued for another 24 h. Sectioning was done on a Leica UC6 ultramicrotome and sections were collected on carbon coated 200 mesh copper grids (Electron Microscopy Sciences, Hatfield, PA). Grids with 90 nm thick sections were placed on a drop of water, sandwiched between two coverslips and mounted in a ring holder (Kukulski et al., 2012). Fluorescence was imaged on the grid with sections facing the objective using an Olympus IX81 inverted widefield microscope equipped with a Hamamatsu ORCA-R2 camera and an Olympus PlApo 100x/1.45 oil objective lens on the same day as the sections had been cut. Subsequently, grids were stained with 3% uranyl acetate and Reynold’s lead citrate and the identical areas, identified by position on the grid in relation to grid center, were imaged on a Tecnai F20 EM (FEI, Eindhoven, The Netherlands) operating at 200kV using the SerialEM software package (Mastronarde, 2005) and an FEI Eagle 4K × 4K CCD camera. Fluorescence and electron microscopy images were correlated based on morphological features using the ICY plug-in ec-CLEM (Paul-Gilloteaux et al., 2017).

### Light microscopy

Imaging of strains expressing Sec63-GFP was done as described (Schuck et al., 2009). Cells were grown to OD_600_ = 0.2 in YPD medium, washed once with SCD medium and resuspended in the same volume of SCD. Cells were left untreated or were treated with 8 mM DTT, grown for 3 h and imaged live at room temperature with a spinning disk confocal microscope consisting of a TE2000U inverted microscope (Nikon, Tokyo, Japan), a CSU22 spinning disk confocal (Yokogawa, Tokyo, Japan), a Cascade II:512 camera (Photometrics, Tucson, Arizona) and a Plan Apo VC 100x/1.4 oil objective lens (Nikon).

For imaging of strains expressing GFP-Pho8 or Hmg2-GFP under the control of the *GAL1* promoter, cells were grown for 10 h in SC-raf medium and diluted into fresh SC-raf to reach early log phase (OD_600_ = 0.1 – 0.5) approximately 16 h later. For induction of the *GAL1* promoter, cells were diluted into SC-raf/gal such that they reached OD_600_ = 2 – 3 after another 24 h. Where indicated, 10 *µ*M CMAC (Thermo Fisher Scientific) or 1 *µ*g/ml FM4-64 (Thermo Fisher Scientific, St. Leon-Rot, Germany) were added for the last 16 h to stain the vacuole lumen or vacuole membrane, respectively. Cells were mounted on a glass coverslip and overlayed with an agarose pad containing SC-raf/gal medium, thus immobilizing the cells and ensuring adequate nutrient supply for at least another 3 h. Images were acquired on the Olympus IX81 widefield microscope at room temperature as described above.

### Flow cytometry

Cells were cultured as above to induce expression of GFP-Pho8. One-hundred *µ*l aliquots were taken to measure GFP fluorescence after excitation with a 488 nm laser with a FACS Canto flow cytometer (BD Biosciences, Franklin Lakes, New Jersey). GFP fluorescence was corrected for cell autofluorescence using a strain not expressing GFP and normalized to cell size using the forward scatter.

### Time-lapse imaging and image analysis

For time-lapse imaging, cells from up to four different strains were mounted on glass coverslips with agarose pads as described above and imaged with a Nikon Ti-E widefield microscope equipped with a motorized stage, the Nikon perfect focus system, a 60x/1.49 oil immersion lens and a Flash4 Hamamatsu sCMOS camera. Movies were taken at a frame rate of one per 2 min over a total imaging period of 2 h. For each frame, two fields of view were imaged for each of the four strains and Z-stacks were acquired that consisted of five optical planes spaced 1 *µ*m apart.

Image analysis was done with the Fiji distribution of ImageJ (Schindelin et al., 2012) extended by plugins and custom code for image classification (available on github). Fluorescent images were subjected to rolling-ball background subtraction with a radius of 80 pixels (8.66 *µ*m). For cell tracking, the CMAC-stained vacuole served as reference. First, image drift over time was corrected in x/y dimensions using the plugin Correct 3D drift with a custom modification by its developer to suppress z-correction. Next, cells were segmented with the Bernsen method for auto local thresholding with radius set to 15 and parameters 1 and 2 set to 0. Images were de-speckled and holes filled in. The resulting binary mask was used as input for the MTrack2 plugin, which was modified such that cropped single-cell movies were generated rather than cell movement tracks. For classification, time-aligned montages of 4×5 cells were generated from single-cell movies that covered frame 30 and at least 5 preceding and 10 following frames. Single-cell movies of all acquired fields of view were randomized during montage generation, enabling blindfolded analysis of different strains. These montages served as input for manual classification by click recording.

To determine the fraction of cells containing a GFP-Pho8 structure (Figure 5A), montages were generated from 100 randomly chosen single cell movies per strain and experiment. Cells were classified as containing or not containing a GFP-Pho8 structure (stack, cytosolic whorl, uptake intermediate and vacuolar whorl) and the fraction of cells with a GFP-Pho8 structure was determined.

For targeted analysis of GFP-Pho8 structures, cells with high GFP-Pho8 levels were selected by creating montages from single-cell movies of cells with GFP intensities between 12,000 and 18,000 arbitrary units. For this, maximum GFP intensities were calculated for all generated single-cell movies within a centered, circular area with diameter covering 70% of the image height and width to exclude neighboring cells from the intensity analysis. Maximum GFP intensities were calculated across all z-slices and time frames of the movies, followed by random selection of at least 200 cells per strain within the set intensity range.

For analysis of the distribution of GFP-Pho8 structures (Figure 5B), the above pre-selected single-cell movies were used to classify cells according to the presence of stacks, cytosolic whorls, uptake intermediates and vacuolar whorls that were present at the start of the movie or arose during imaging. Multiple assignments per cell were allowed. If transitions between classes of structures occurred during a movie, the structure in question was assigned to the latest observed step in the process, with the order from early to late being stack to uptake intermediate to vacuolar whorl or stack to cytosolic whorl. The fraction of each GFP-Pho8 structure relative to the total number of GFP-Pho8 structures was determined.

For analysis of the localization of stacks (Figure 5C), the above pre-selected single-cell movies were used to classify cells according to stack type (vacuole-associated, perinuclear or peripheral). Multiple assignments per cell were allowed. The fractions of vacuole-associated, perinuclear and peripheral stacks relative to the total number of stacks were determined.

For analysis of uptake intermediate fate (Figure 2C), cells annotated as uptake intermediates or vacuolar whorls in the analysis above were used as input. Single-cell movies showing transitions of uptake intermediates to vacuolar whorls or stacks were counted, uptake intermediates showing no transition were ignored. The fractions of intermediate-to-whorl and intermediate-to-stack conversions relative to the total number of events were determined. The time for conversion of bent stacks to vacuolar whorls (Figure S2A) was determined from cells showing complete stack-to-intermediate-to-whorl transitions.

### Microscopy-based genetic screen

Using strain SSY1653, mCherry-tagged *SEC63* and four copies of the *P_GAL1_-GFP-PHO8* expression cassette were introduced into the yeast knockout collection (Open Biosystems, provided by Michael Knop, ZMBH) by synthetic genetic array methodology (Tong and Boone, 2007). For high-troughput microscopy, cells were grown to saturation in 150 *µ*l SC-raf/gal in 96-well plates. New 1 ml cultures in SC-raf/gal in 96-deep-well plates were inoculated with 20 *µ*l saturated cultures and grown overnight. 100 *µ*l of the overnight cultures were transferred into new 96-well plates. Cells were treated with 0.2 *µ*g/ml rapamycin (Sigma) to increase the fraction of cells with a single, fused vacuole and vacuoles were stained with 50 *µ*M CMAC for 5 h. Five *µ*l of the cultures were diluted in 90 *µ*l SC-raf/gal containing low fluorescence yeast nitrogen base without amino acids, folic acid and riboflavin (Formedium, Norfolk, UK) and 0.2 *µ*g/ml rapamycin in 96-well glass bottom plates (Brooks Life Sciences, Chelmsford, Massachusetts) coated with concanavalin A, type IV (Sigma). After 1 h of incubation, non-attached cells were removed by aspiration of the supernatant and addition of 180 *µ*l fresh SC-raf/gal containing low fluorescence yeast nitrogen base and rapamycin. The strain collection was imaged using the Nikon Ti-E widefield microscope described above. Z-stacks with 5 planes of 1 *µ*m width were acquired at three fields of view per mutant.

To facilitate subsequent image analysis, automated cell segmentation was performed. Images were loaded into MATLAB and background fluorescence was locally removed by applying a morphological opening to the images and subtracting the result from the original images. Cells were automatically segmented based on the bright field images. The Z-stack with the highest contrast was selected for further processing based on the standard deviation of pixel intensities in the entire image. The coordinates of initial cell objects were identified using a circular Hough transformation and searching for objects with a radius range of 15 - 25 pixels (1.6 - 2.7 *µ*m). A bounding box extending 2.7 *µ*m beyond these objects was then cropped and each cell was further processed individually. A strong background correction was applied to the crops by subtracting 1.5 times the local background intensity as determined above. Images were sharpened by unsharp masking and canny edge detection was applied to create a binary image. A best fit ellipse was calculated by applying a modified version of the algorithm ‘ellipse detection using 1D Hough transform’ by Martin Simonovsky (downloaded from MATLAB file exchange February 16, 2016). Vacuoles were further segmented using the CMAC image. A 3×3 median smoothing filter was applied and CMAC staining was locally defined as pixels brighter than the average pixel intensity of a surrounding ring of radius equal to 9 pixels (= 1 *µ*m). If these pixels were brighter than an empirically defined background threshold, they were designated vacuolar. Objects smaller than 10 pixels in area were removed as noise and segmented vacuole objects were smoothed by morphological closing. Basic cell measurements were then based on these segmented objects and included total cell GFP and RFP intensity across all Z-stacks, and total vacuole GFP and RFP intensity as segmented per Z-stack.

Using custom MATLAB viewer software, montages were generated containing 90 uniformly arrayed cells per field of view based on the identified cell crops. These montages were more conducive to visual inspection than the original images with a random spread of cells. Images were analyzed visually to identify mutants in which the frequency of GFP-Pho8 stretches and rings clearly differed from the wild-type.

### Pho8 assay

Cells were grown to mid log phase (OD_600_ = 0.1 – 0.5) in YPD medium. For measurement of activity in untreated cells, cultures were diluted to OD_600_ = 0.4 in YPD and grown for 2 h. For measurement of activity in tunicamycin-treated cells, cultures were diluted to OD_600_ = 0.4 in YPD, treated with 1 *µ*g/ml tunicamycin (Merck) and grown for 9 h. For measurement of activity in nitrogen-starved cells, cells were diluted to OD_600_ = 0.4 in YPD, pelleted by centrifugation at 1,500 xg for 5 min, resuspended in water, pelleted again, resuspended in the same volume of SCD-N medium (0.7% yeast nitrogen base without ammonium sulfate from Formedium, Norfolk, UK, and 2% glucose) and grown for 6 h. *Δpep4* strains were additionally treated with 1 mM PMSF to block Pho8Δ60 reporter activation. Cells were collected by centrifugation, washed with water, resuspended in 200 *µ*l cold lysis buffer (20 mM PIPES pH 6.8, 1% Triton X-100, 50 mM KCl, 100 mM KOAc pH 7.5, 10 mM MgSO_4_, 10 *µ*M ZnSO_4_, 1 mM PMSF and complete protease inhibitors from Roche), and disrupted by bead beating with a FastPrep 24 (MP Biomedical, Heidelberg, Germany). Lysates were adjusted to 1, 2 or 4 ODs/100 *µ*l (endogenous Pho8, cyto-Pho8Δ60 or Yop1-Pho8Δ60, respectively) in lysis buffer and cleared by centrifugation at 11,000 xg for 2 min at 4°C. For each reaction, 50 *µ*l lysate were combined with 200 *µ*l reaction buffer (250 mM Tris pH 8.5, 1% Triton X-100, 10 mM MgSO_4_, 10 *µ*M ZnSO_4_) containing 1.25 mM p-nitrophenyl phosphate (pNPP, Sigma) as substrate and incubated at 37°C for 5 – 20 min. Reactions were stopped by addition of 250 *µ*l 1 M glycine/KOH pH 11. Absorption was measured at 405 nm. To correct for background absorption arising from buffer components, lysate components or non-specific pNPP hydrolysis, the absorption of a buffer control (containing no pNPP and no cell lysate), a lysate control (containing no pNPP) and a substrate control (containing no cell lysate) were determined and subtracted from the absorption of the experimental samples. Protein concentration of cell lysates was determined with the BCA assay kit (Thermo Fisher Scientific, Schwerte, Germany).

One unit of specific phosphatase activity was defined as 1 nmol p-nitrophenol produced per min and mg total protein. To account for background phosphatase activity in strains containing Yop1-Pho8Δ60 and cyto-Pho8Δ60, the activity in *Δpep4* cells was subtracted from the phosphatase activity in all other samples. Similarly, for measurement of endogenous Pho8, the activity in *Δpho8* cells was subtracted from the phosphatase activity in all other samples. The remaining activity was expressed as % of the activity in wild-type cells.

---

**Figure S1.**
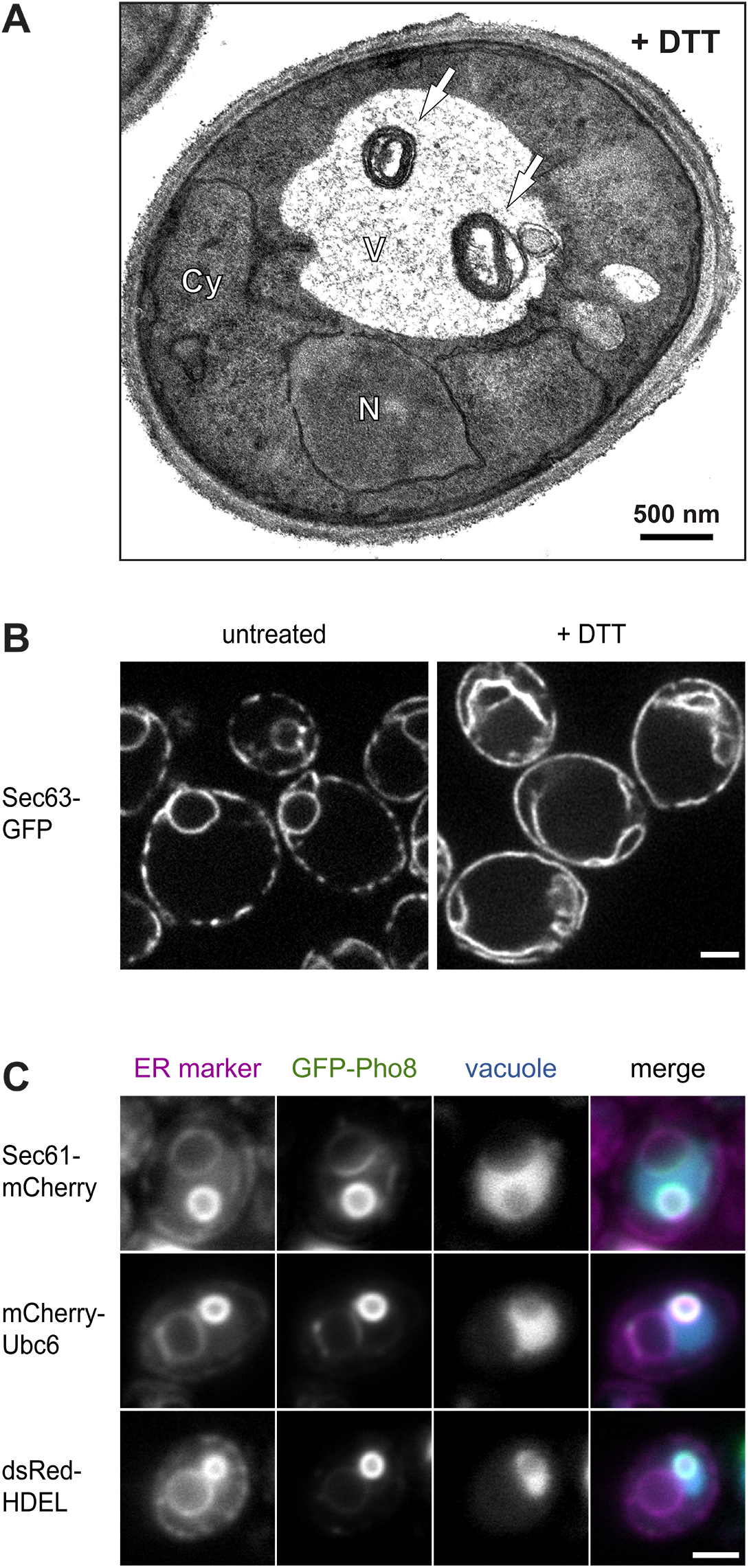
Expression of ER-resident transmembrane proteins can induce autophagy of ER whorls. **(A)** Electron micrograph of a yeast cell treated with DTT for 3 h. White arrows indicate ER whorls in the vacuole. Cy, cytosol; N, nucleus; V, vacuole. **(B)** Fluorescence images of cells expressing the general ER marker Sec63-GFP. Cells were left untreated or treated with DTT for 3 h. DTT induces expansion of the peripheral ER and the formation of ER extensions into the cytosol, but no structures resembling whorls are visible. Scale bar: 2 *µ*m. **(C)** Fluorescence images of CMAC-stained cells expressing GFP-Pho8 and the ER markers Sec61-mCherry, mCherry-Ubc6 or dsRed-HDEL. GFP-Pho8 and the ER markers highlight ring structures inside CMAC-stained vacuoles. Scale bar: 2 *µ*m.

**Figure S2.**
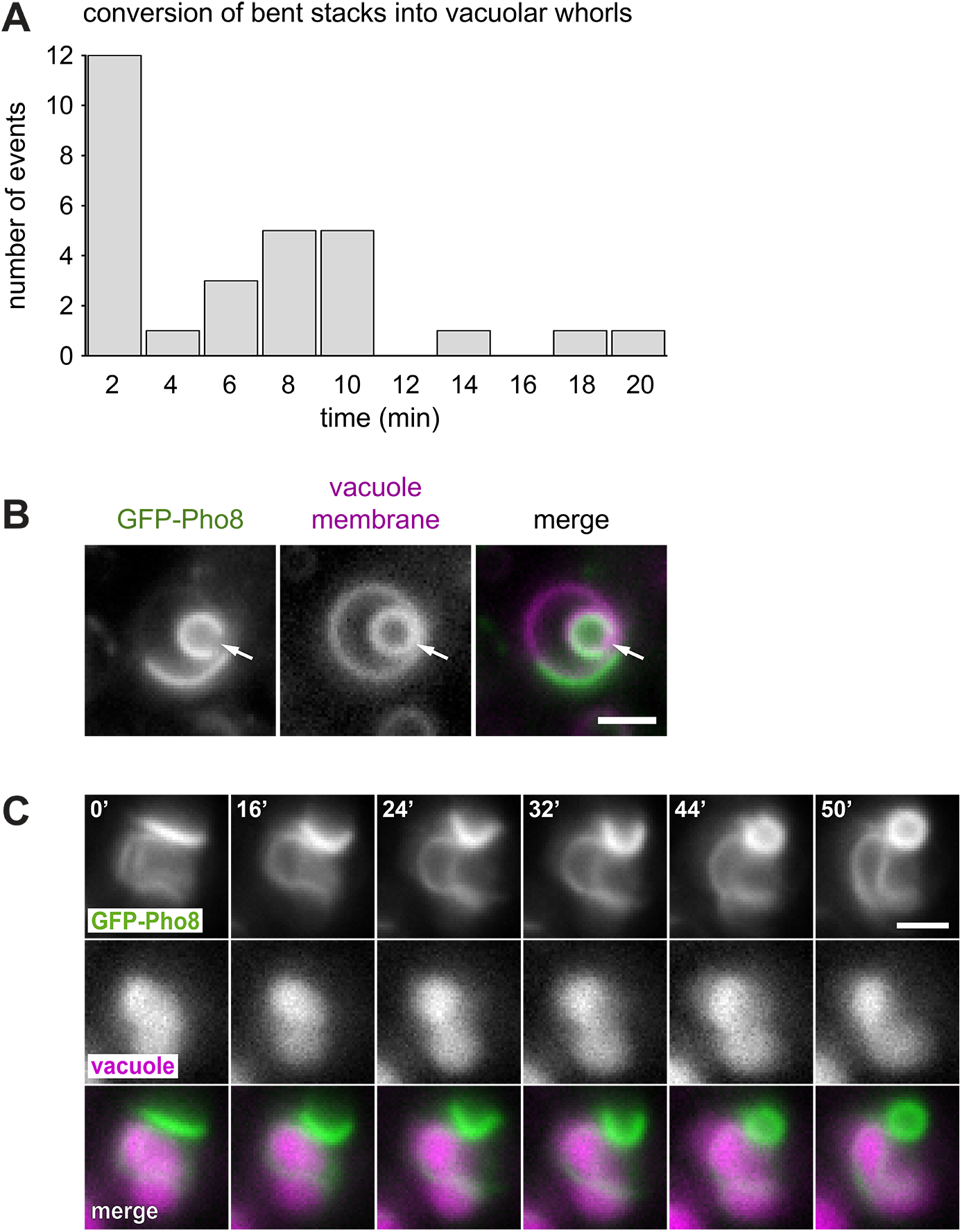
Autophagy of ER whorls occurs by microautophagy. **(A)** Conversion times of bent stacks into vacuolar whorls. n = 29. **(B)** Fluorescence images of a cell expressing GFP-Pho8 and stained with the vacuole membrane dye FM4-64. The whorl is completely surrounded by vacuole membrane but has an opening (arrow). Scale bar: 2 *µ*m. **(C)** Individual frames from time-lapse imaging of a cell expressing GFP-Pho8 and stained with the vacuole dye CMAC. The image sequence shows the conversion of a peripheral stack into a cytosolic whorl. Numbers indicate the time in minutes after the start of the image sequence. Scale bar: 2 *µ*m.

**Figure S3.**
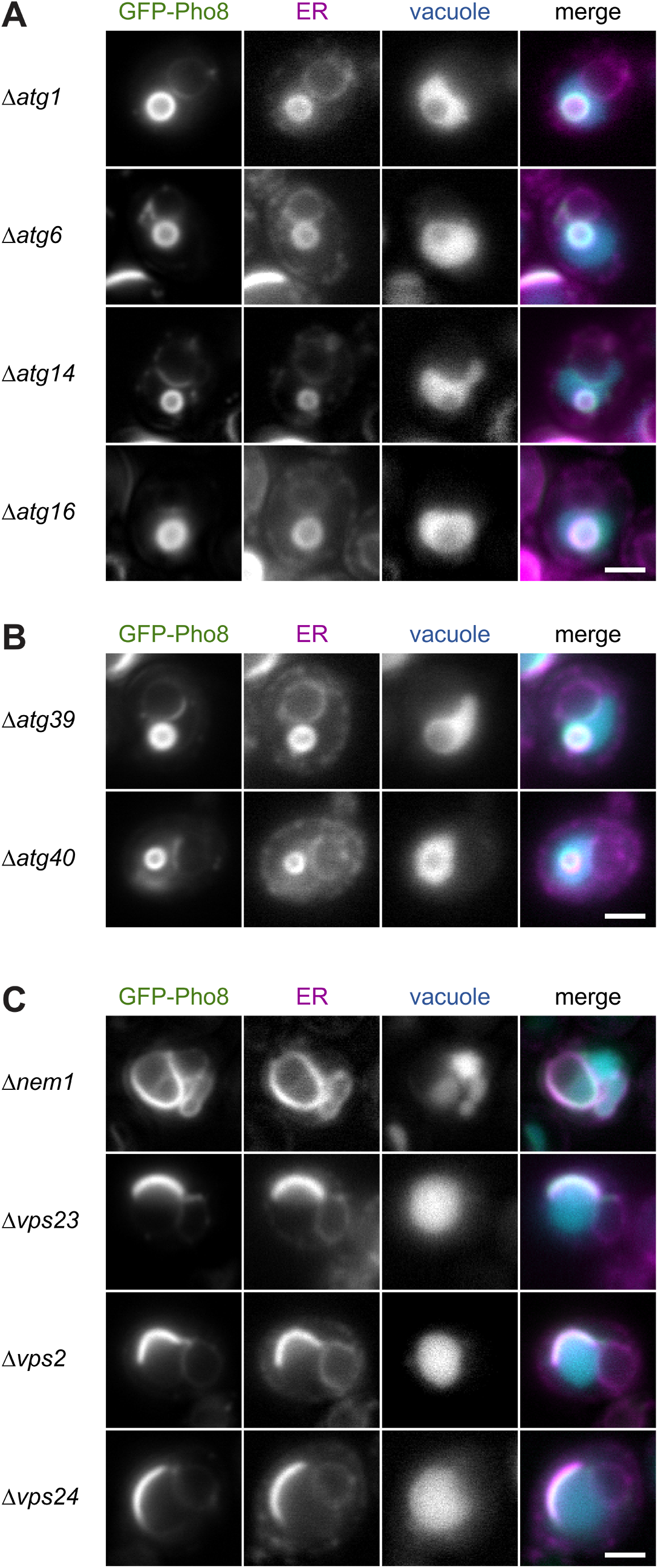
The Nem1-Spo7 complex and ESCRT proteins are key components for micro-ER-phagy. **(A)** Fluorescence images of *Δatg1*, *Δatg6*, *Δatg14* and *Δatg16* cells expressing GFP-Pho8 and the general ER marker Sec63-mCherry and stained with CMAC to label the vacuole. Scale bar: 2 *µ*m. **(B)** As in panel A but of *Δatg39* and *Δatg40* cells. **(C)** As in panel A but of *Δnem1*, *Δvps23*, *Δvps2* and *Δvps24* cells. Cells lacking Nem1 have irregularly shaped nuclei and expanded perinuclear as well as peripheral ER (Santos-Rosa et al., 2005).

**Figure S4.**
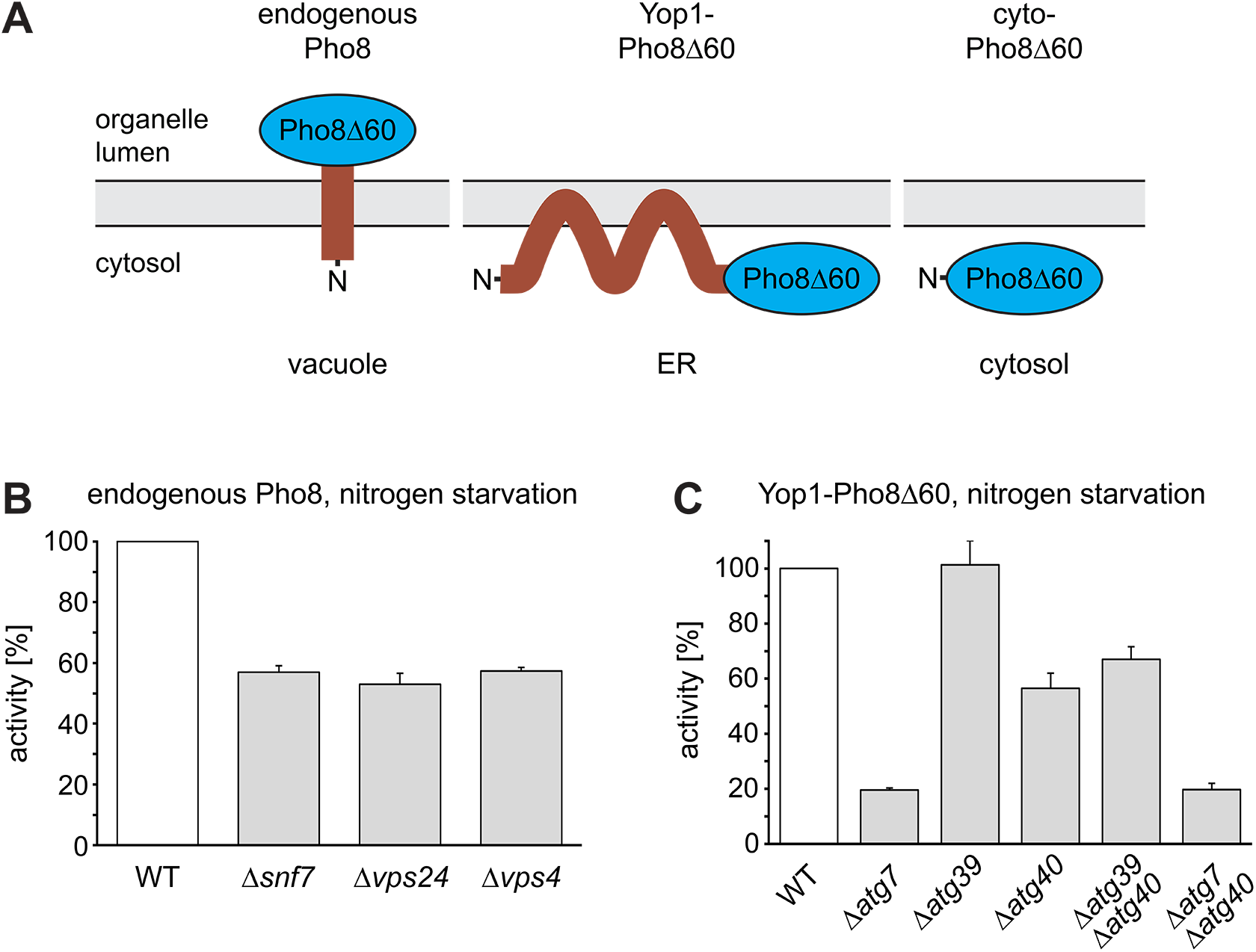
Macro- and micro-ER-phagy are parallel pathways with distinct molecular requirements. **(A)** Pho8-based autophagy reporters. **(B)** Relative activity of endogenous Pho8 after nitrogen starvation. Mean ± SEM, n = 3. **(C)** Relative activity of the ER-phagy reporter Yop1-Pho8Δ60 after nitrogen starvation. *ATG40* is involved in autophagy of Yop1-Pho8Δ60 and is epistatic with *ATG7*. Mean ± SEM, n ≥ 3.

**Figure S5.**
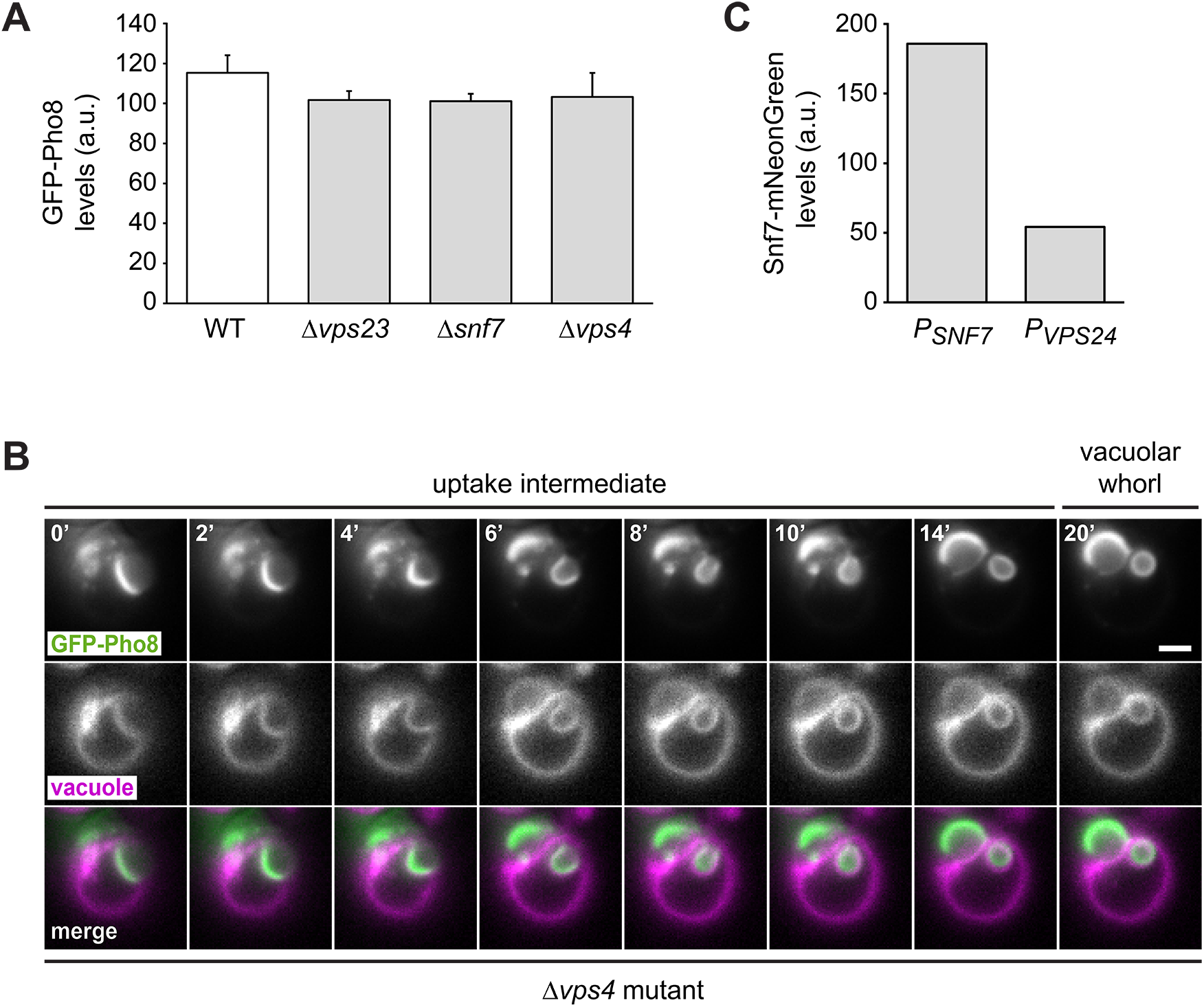
ESCRT machinery mediates vacuole membrane dynamics during micro-ER-phagy of GFP-Pho8 structures. **(A)** GFP-Pho8 expression levels as measured by flow cytometry. Mean ± SEM, n ≥ 3. **(B)** Individual frames from time-lapse imaging of a *Δvps4* cell expressing GFP-Pho8 and stained with the vacuole membrane dye FM4-64. Numbers indicate the time in minutes after the start of the image sequence. Scale bar: 2 *µ*m. **(C)** Snf7-mNeonGreen levels as measured by flow cytometry upon expression from the *SNF7* or *VPS24* promoter.

**Movie 1.** Snf7 transiently associates with sites of ER whorl uptake into the vacuole. Movie sequence of a cell expressing non-fluorescent GFP(Y66F)-Pho8, Snf7-mNeonGreen and the ER marker mCherry-Ubc6 and stained with the vacuole dye CMAC. The movie contains 30 frames (t1 - t30) acquired at 2-minute intervals. Each frame consists of five optical sections spaced 1 *µ*m apart (z1 - z5). The movie sequence starts with a vacuole-associated stack (t1, z1) that bends, forms an uptake intermediate (t4, z1; 0’ in Figure 5D) and finally a vacuolar whorl (t6, z2; 4’ in Figure 5D). At these stages, no Snf7 is apparent in the vicinity of the whorl in any of the optical sections. In the next frame, a Snf7 punctum appears next to the rim of the whorl (t7, z1; 6’ in Figure 5D), showing recruitment of Snf7 within 2 minutes or less. The Snf7 punctum remains adjacent to the whorl for 15 frames (30 minutes). During this time, the punctum is either at the side (e.g. t13, z1) or the top of the whorl (t15, compare z1 - z4), indicating that Snf7 associates with the rim rather than the lumen of the whorl and does not represent an endosome trapped inside the whorl. Starting at frame 19, the Snf7 punctum and the whorl are present in the same focal plane (t19, z1; 30’ in Figure 5D). Optical sections in which a vacuole membrane invagination is visible show that the whorl-associated Snf7 punctum is oriented towards the cytosolic opening of the invagination and immediately adjacent to the vacuole membrane (t22, z2; 36’ in Figure 5D). This suggests that Snf7 is located at the neck of the invagination. The Snf7 signal drops in the next frame (t23, z1) and completely disappears in the one after that, so that the vacuolar whorl is free of Snf7 (t24, z1 - z5; 40’ in Figure 5D). Other Snf7 puncta in the same and neighbouring cells remain during these frames, indicating that the disappearance of the whorl-associated Snf7 punctum is not due to photobleaching.

**Movie 2.** Snf7 puncta at ER whorls are oriented towards the neck of vacuole membrane invaginations. Movie sequence of a cell expressing non-fluorescent GFP(Y66F)-Pho8, Snf7-mNeonGreen and the ER marker mCherry-Ubc6 and stained with the vacuole dye CMAC. The movie contains 12 frames (t1 – t12) acquired at 2-minute intervals. Each frame consists of five optical sections spaced 1 *µ*m apart (z1 - z5). The movie sequence starts with a vacuolar whorl in a vacuole membrane invagination and a Snf7 punctum that is positioned at the rim of the vacuolar whorl (t1, z4; 0’ in Figure 5F). The Snf7 punctum is at the neck of the invaginated vacuole membrane, where it remains throughout the movie. Optical section t4, z4 corresponds to 6’ in Figure 5F, section t7, z4 to 12’ and section t12, z4 to 22’. Note that the three channels are shifted against each other at t11 and that the intensity of the Snf7-mNeonGreen signal decays over the course of the movie.

**Table S2.** Plasmids used in this study.

| Plasmid | Alias | Source |
| --- | --- | --- |
| pRS406-P<sub>GPD</sub>-GFP-Pho8 | pSS503 | this study |
| pRS406-P<sub>GPD</sub>-GFP(L221K)-Pho8 | pSS611 | this study |
| pRS416-P<sub>GAL</sub> | pSS031 | Mumberg et al., 1994 |
| pRS406-P<sub>GAL</sub>-GFP-Pho8 | pSS590 | this study |
| pRS415 | pSS008 | Sikorski and Hieter, 1989 |
| pRS405-P<sub>GAL</sub>-GFP-Pho8 | pSS591 | this study |
| Yip-P<sub>GAL</sub>-Hmg2(K6R)-GFP | pRH1767 | Federovitch et al., 2008 |
| pRS415-P<sub>ADH</sub> | pSS022 | Mumberg et al., 1995 |
| pRS415-P<sub>ADH</sub>-Hmg2(K6R)-GFP | pSS327 | this study |
| pRS406-P<sub>GAL</sub>-Hmg2(K6R)-GFP | pSS955 | this study |
| Yiplac-ssDsRed-HDEL-nat | pKW1803 | Madrid et al., 2006 |
| pRS406-P<sub>GPD</sub>-mCherry-Ubc6 | pSS117 | Benoit Kornmann |
| pRS404 | pSS003 | Sikorski and Hieter, 1989 |
| pRS404-P<sub>GPD</sub>-mCherry-Ubc6 | pSS962 | this study |
| pRS416-Snf7-LAP-eGFP | pSS867 | Adell et al., 2017 |
| pFA6a-mNeonGreen-kanMX6 | pSS445 | this study |
| pRS416-P<sub>SNF7</sub>-Snf7-LAP-mNeonGreen | pSS931 | this study |
| pRS416-P<sub>VPS24</sub>-Snf7-LAP-mNeonGreen | pSS951 | this study |
| pRS405-P<sub>GAL</sub>-GFP(Y66F)Pho8 | pSS915 | this study |

**Table S3.**
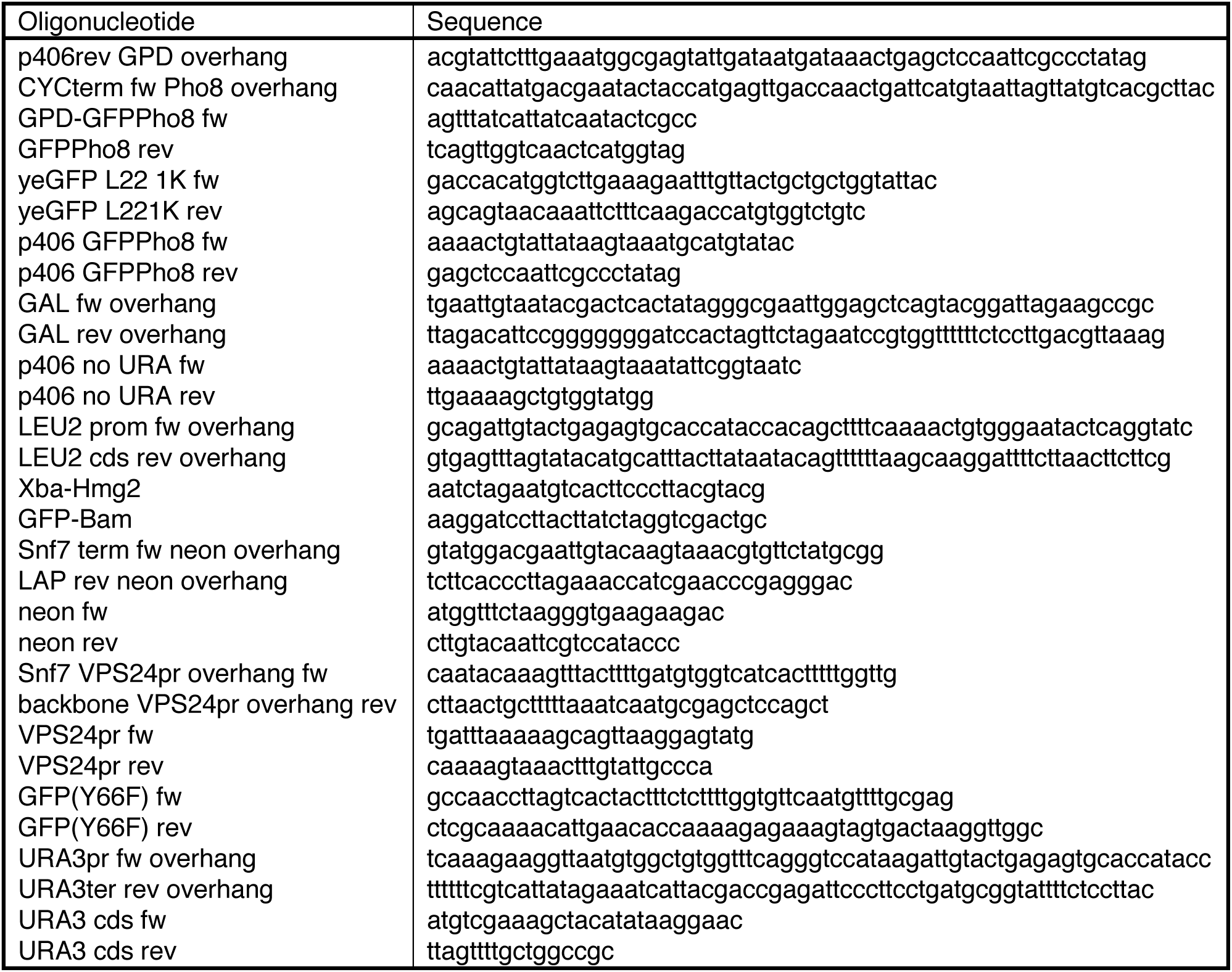
Oligonucleotides used in this study.

**Table S4.**
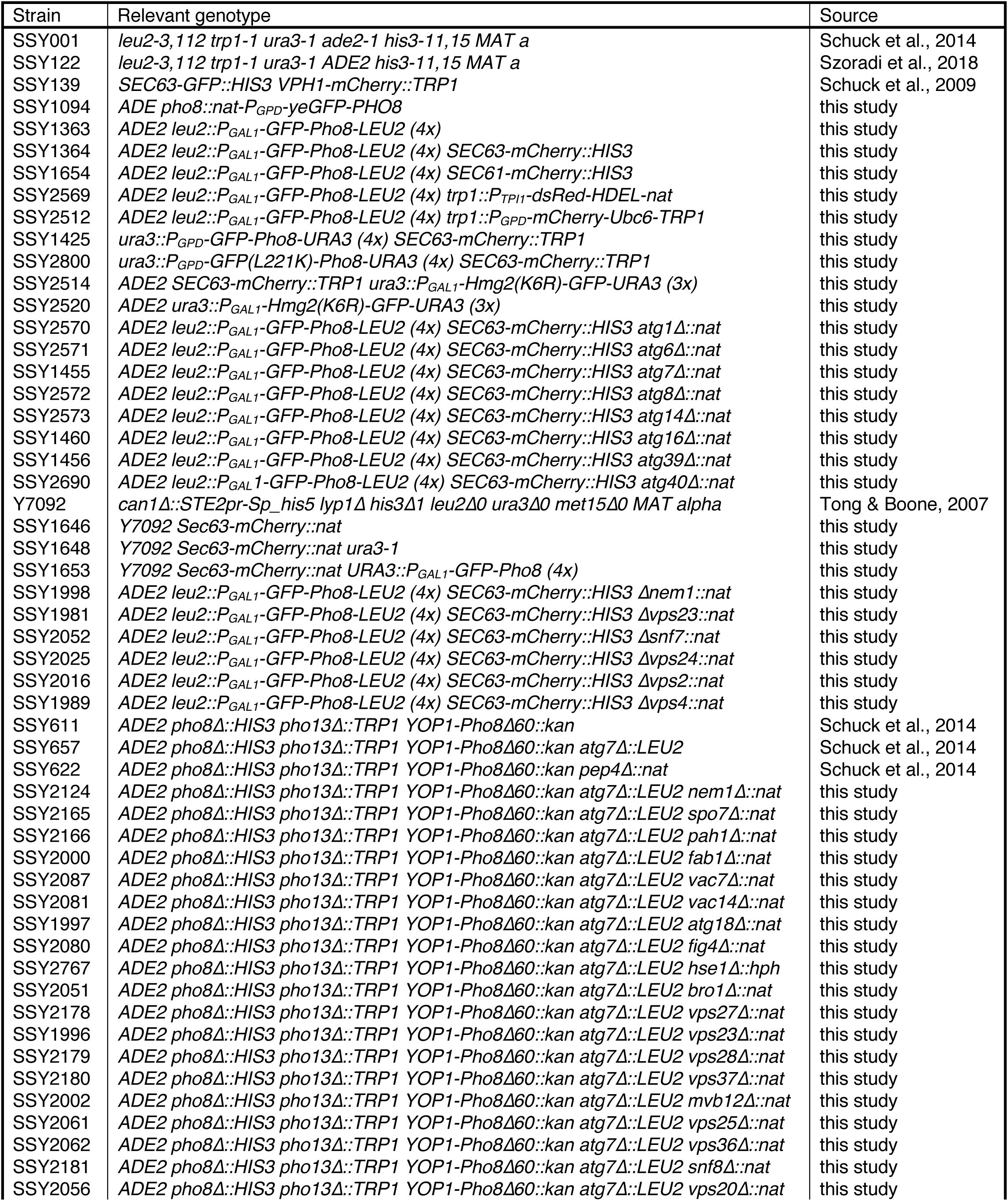
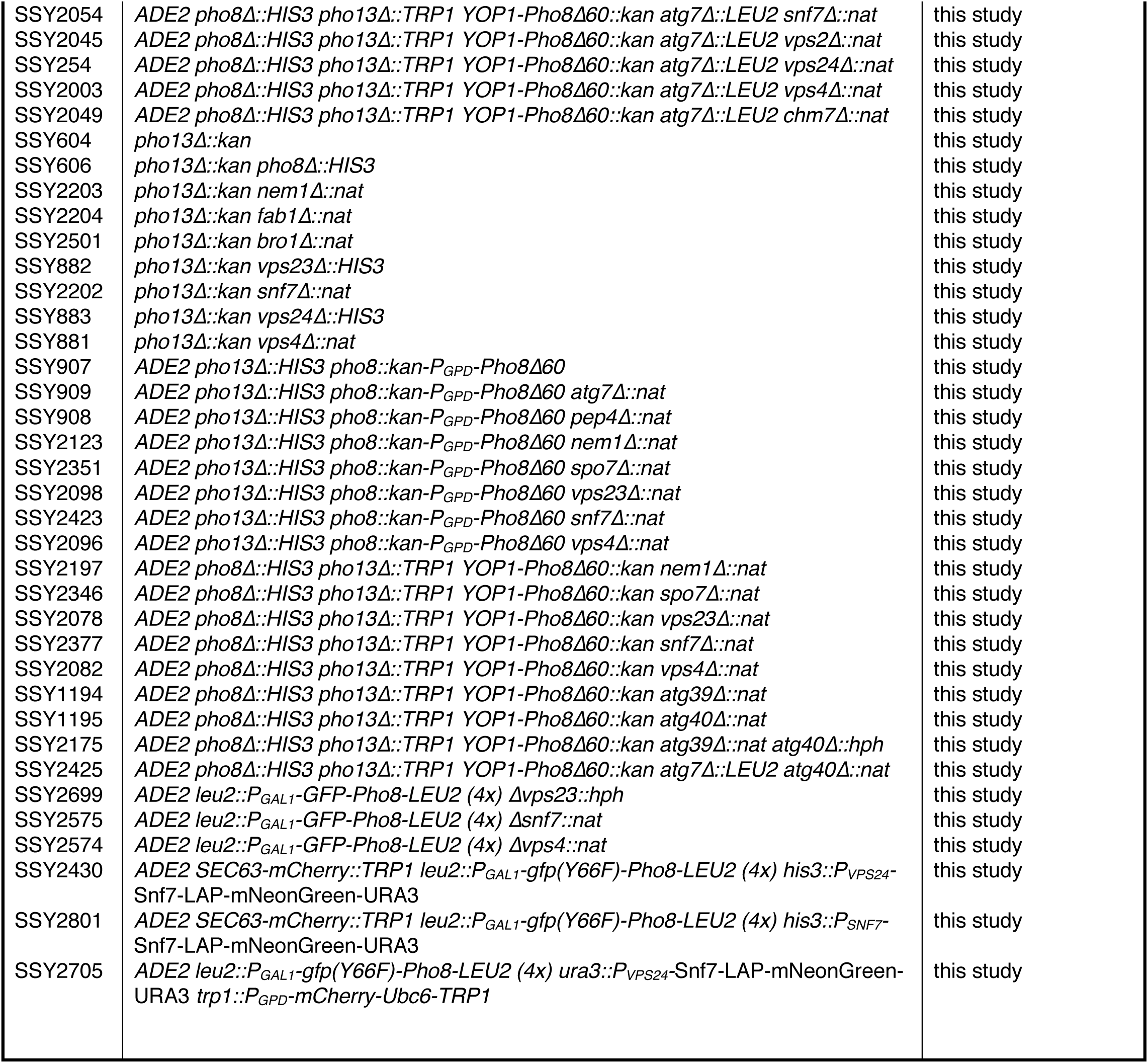
Yeast strains used in this study.

| Strain | Relevant genotype | Source |
| --- | --- | --- |
| SSY001 | *leu2-3,112 trp1-1 ura3-1 ade2-1 his3-11,15 MAT a* | Schuck et al., 2014 |
| SSY122 | *leu2-3,112 trp1-1 ura3-1 ADE2 his3-11,15 MAT a* | Szoradi et al., 2018 |
| SSY139 | *SEC63-GFP::HIS3 VPH1-mCherry::TRP1* | Schuck et al., 2009 |
| SSY1094 | *ADE2 pho8::nat-P<sub>GPD</sub>-yeGFP-PHO8* | this study |
| SSY1363 | *ADE2 leu2::P<sub>GAL1</sub>-GFP-Pho8-LEU2 (4x)* | this study |
| SSY1364 | *ADE2 leu2::P<sub>GAL1</sub>-GFP-Pho8-LEU2 (4x) SEC63-mCherry::HIS3* | this study |
| SSY1654 | *ADE2 leu2::P<sub>GAL1</sub>-GFP-Pho8-LEU2 (4x) SEC61-mCherry::HIS3* | this study |
| SSY2569 | *ADE2 leu2::P<sub>GAL1</sub>-GFP-Pho8-LEU2 (4x) trp1::P<sub>TPI1</sub>-dsRed-HDEL-nat* | this study |
| SSY2512 | *ADE2 leu2::P<sub>GAL1</sub>-GFP-Pho8-LEU2 (4x) trp1::P<sub>GPD</sub>-mCherry-Ubc6-TRP1* | this study |
| SSY1425 | *ura3::P<sub>GPD</sub>-GFP-Pho8-URA3 (4x) SEC63-mCherry::TRP1* | this study |
| SSY2800 | *ura3::P<sub>GPD</sub>-GFP(L221K)-Pho8-URA3 (4x) SEC63-mCherry::TRP1* | this study |
| SSY2514 | *ADE2 SEC63-mCherry::TRP1 ura3::P<sub>GAL1</sub>-Hmg2(K6R)-GFP-URA3 (3x)* | this study |
| SSY2520 | *ADE2 ura3::P<sub>GAL1</sub>-Hmg2(K6R)-GFP-URA3 (3x)* | this study |
| SSY2570 | *ADE2 leu2::P<sub>GAL1</sub>-GFP-Pho8-LEU2 (4x) SEC63-mCherry::HIS3 atg1Δ::nat* | this study |
| SSY2571 | *ADE2 leu2::P<sub>GAL1</sub>-GFP-Pho8-LEU2 (4x) SEC63-mCherry::HIS3 atg6Δ::nat* | this study |
| SSY1455 | *ADE2 leu2::P<sub>GAL1</sub>-GFP-Pho8-LEU2 (4x) SEC63-mCherry::HIS3 atg7Δ::nat* | this study |
| SSY2572 | *ADE2 leu2::P<sub>GAL1</sub>-GFP-Pho8-LEU2 (4x) SEC63-mCherry::HIS3 atg8Δ::nat* | this study |
| SSY2573 | *ADE2 leu2::P<sub>GAL1</sub>-GFP-Pho8-LEU2 (4x) SEC63-mCherry::HIS3 atg14Δ::nat* | this study |
| SSY1460 | *ADE2 leu2::P<sub>GAL1</sub>-GFP-Pho8-LEU2 (4x) SEC63-mCherry::HIS3 atg16Δ::nat* | this study |
| SSY1456 | *ADE2 leu2::P<sub>GAL1</sub>-GFP-Pho8-LEU2 (4x) SEC63-mCherry::HIS3 atg39Δ::nat* | this study |
| SSY2690 | *ADE2 leu2::P<sub>GAL1</sub>-GFP-Pho8-LEU2 (4x) SEC63-mCherry::HIS3 atg40Δ::nat* | this study |
| Y7092 | *can1Δ::STE2pr-Sp_his5 lyp1Δ his3Δ1 leu2Δ0 ura3Δ0 met15Δ0 MAT alpha* | Tong & Boone, 2007 |
| SSY1646 | *Y7092 Sec63-mCherry::nat* | this study |
| SSY1648 | *Y7092 Sec63-mCherry::nat ura3-1* | this study |
| SSY1653 | *Y7092 Sec63-mCherry::nat URA3::P<sub>GAL1</sub>-GFP-Pho8 (4x)* | this study |
| SSY1998 | *ADE2 leu2::P<sub>GAL1</sub>-GFP-Pho8-LEU2 (4x) SEC63-mCherry::HIS3 Δnem1::nat* | this study |
| SSY1981 | *ADE2 leu2::P<sub>GAL1</sub>-GFP-Pho8-LEU2 (4x) SEC63-mCherry::HIS3 Δvps23::nat* | this study |
| SSY2052 | *ADE2 leu2::P<sub>GAL1</sub>-GFP-Pho8-LEU2 (4x) SEC63-mCherry::HIS3 Δsnf7::nat* | this study |
| SSY2025 | *ADE2 leu2::P<sub>GAL1</sub>-GFP-Pho8-LEU2 (4x) SEC63-mCherry::HIS3 Δvps24::nat* | this study |
| SSY2016 | *ADE2 leu2::P<sub>GAL1</sub>-GFP-Pho8-LEU2 (4x) SEC63-mCherry::HIS3 Δvps2::nat* | this study |
| SSY1989 | *ADE2 leu2::P<sub>GAL1</sub>-GFP-Pho8-LEU2 (4x) SEC63-mCherry::HIS3 Δvps4::nat* | this study |
| SSY611 | *ADE2 pho8Δ::HIS3 pho13Δ::TRP1 YOP1-Pho8Δ60::kan* | Schuck et al., 2014 |
| SSY657 | *ADE2 pho8Δ::HIS3 pho13Δ::TRP1 YOP1-Pho8Δ60::kan atg7Δ::LEU2* | Schuck et al., 2014 |
| SSY622 | *ADE2 pho8Δ::HIS3 pho13Δ::TRP1 YOP1-Pho8Δ60::kan pep4Δ::nat* | Schuck et al., 2014 |
| SSY2124 | *ADE2 pho8Δ::HIS3 pho13Δ::TRP1 YOP1-Pho8Δ60::kan atg7Δ::LEU2 nem1Δ::nat* | this study |
| SSY2165 | *ADE2 pho8Δ::HIS3 pho13Δ::TRP1 YOP1-Pho8Δ60::kan atg7Δ::LEU2 spo7Δ::nat* | this study |
| SSY2166 | *ADE2 pho8Δ::HIS3 pho13Δ::TRP1 YOP1-Pho8Δ60::kan atg7Δ::LEU2 pah1Δ::nat* | this study |
| SSY2000 | *ADE2 pho8Δ::HIS3 pho13Δ::TRP1 YOP1-Pho8Δ60::kan atg7Δ::LEU2 fab1Δ::nat* | this study |
| SSY2087 | *ADE2 pho8Δ::HIS3 pho13Δ::TRP1 YOP1-Pho8Δ60::kan atg7Δ::LEU2 vac7Δ::nat* | this study |
| SSY2081 | *ADE2 pho8Δ::HIS3 pho13Δ::TRP1 YOP1-Pho8Δ60::kan atg7Δ::LEU2 vac14Δ::nat* | this study |
| SSY1997 | *ADE2 pho8Δ::HIS3 pho13Δ::TRP1 YOP1-Pho8Δ60::kan atg7Δ::LEU2 atg18Δ::nat* | this study |
| SSY2080 | *ADE2 pho8Δ::HIS3 pho13Δ::TRP1 YOP1-Pho8Δ60::kan atg7Δ::LEU2 fig4Δ::nat* | this study |
| SSY2767 | *ADE2 pho8Δ::HIS3 pho13Δ::TRP1 YOP1-Pho8Δ60::kan atg7Δ::LEU2 hse1Δ::hph* | this study |
| SSY2051 | *ADE2 pho8Δ::HIS3 pho13Δ::TRP1 YOP1-Pho8Δ60::kan atg7Δ::LEU2 bro1Δ::nat* | this study |
| SSY2178 | *ADE2 pho8Δ::HIS3 pho13Δ::TRP1 YOP1-Pho8Δ60::kan atg7Δ::LEU2 vps27Δ::nat* | this study |
| SSY1996 | *ADE2 pho8Δ::HIS3 pho13Δ::TRP1 YOP1-Pho8Δ60::kan atg7Δ::LEU2 vps23Δ::nat* | this study |
| SSY2179 | *ADE2 pho8Δ::HIS3 pho13Δ::TRP1 YOP1-Pho8Δ60::kan atg7Δ::LEU2 vps28Δ::nat* | this study |
| SSY2180 | *ADE2 pho8Δ::HIS3 pho13Δ::TRP1 YOP1-Pho8Δ60::kan atg7Δ::LEU2 vps37Δ::nat* | this study |
| SSY2002 | *ADE2 pho8Δ::HIS3 pho13Δ::TRP1 YOP1-Pho8Δ60::kan atg7Δ::LEU2 mvb12Δ::nat* | this study |
| SSY2061 | *ADE2 pho8Δ::HIS3 pho13Δ::TRP1 YOP1-Pho8Δ60::kan atg7Δ::LEU2 vps25Δ::nat* | this study |
| SSY2062 | *ADE2 pho8Δ::HIS3 pho13Δ::TRP1 YOP1-Pho8Δ60::kan atg7Δ::LEU2 vps36Δ::nat* | this study |
| SSY2181 | *ADE2 pho8Δ::HIS3 pho13Δ::TRP1 YOP1-Pho8Δ60::kan atg7Δ::LEU2 snf8Δ::nat* | this study |
| SSY2056 | *ADE2 pho8Δ::HIS3 pho13Δ::TRP1 YOP1-Pho8Δ60::kan atg7Δ::LEU2 vps20Δ::nat* | this study |
| SSY2054 | ADE2 *pho8Δ::HIS3 pho13Δ::TRP1 YOP1-Pho8Δ60::kan atg7Δ::LEU2 snf7Δ::nat* | this study |
| SSY2045 | ADE2 *pho8Δ::HIS3 pho13Δ::TRP1 YOP1-Pho8Δ60::kan atg7Δ::LEU2 vps2Δ::nat* | this study |
| SSY254 | ADE2 *pho8Δ::HIS3 pho13Δ::TRP1 YOP1-Pho8Δ60::kan atg7Δ::LEU2 vps24Δ::nat* | this study |
| SSY2003 | ADE2 *pho8Δ::HIS3 pho13Δ::TRP1 YOP1-Pho8Δ60::kan atg7Δ::LEU2 vps4Δ::nat* | this study |
| SSY2049 | ADE2 *pho8Δ::HIS3 pho13Δ::TRP1 YOP1-Pho8Δ60::kan atg7Δ::LEU2 chm7Δ::nat* | this study |
| SSY604 | *pho13Δ::kan* | this study |
| SSY606 | *pho13Δ::kan pho8Δ::HIS3* | this study |
| SSY2203 | *pho13Δ::kan nem1Δ::nat* | this study |
| SSY2204 | *pho13Δ::kan fab1Δ::nat* | this study |
| SSY2501 | *pho13Δ::kan bro1Δ::nat* | this study |
| SSY882 | *pho13Δ::kan vps23Δ::HIS3* | this study |
| SSY2202 | *pho13Δ::kan snf7Δ::nat* | this study |
| SSY883 | *pho13Δ::kan vps24Δ::HIS3* | this study |
| SSY881 | *pho13Δ::kan vps4Δ::nat* | this study |
| SSY907 | ADE2 *pho13Δ::HIS3 pho8::kan-P<sub>GPD</sub>-Pho8Δ60* | this study |
| SSY909 | ADE2 *pho13Δ::HIS3 pho8::kan-P<sub>GPD</sub>-Pho8Δ60 atg7Δ::nat* | this study |
| SSY908 | ADE2 *pho13Δ::HIS3 pho8::kan-P<sub>GPD</sub>-Pho8Δ60 pep4Δ::nat* | this study |
| SSY2123 | ADE2 *pho13Δ::HIS3 pho8::kan-P<sub>GPD</sub>-Pho8Δ60 nem1Δ::nat* | this study |
| SSY2351 | ADE2 *pho13Δ::HIS3 pho8::kan-P<sub>GPD</sub>-Pho8Δ60 spo7Δ::nat* | this study |
| SSY2098 | ADE2 *pho13Δ::HIS3 pho8::kan-P<sub>GPD</sub>-Pho8Δ60 vps23Δ::nat* | this study |
| SSY2423 | ADE2 *pho13Δ::HIS3 pho8::kan-P<sub>GPD</sub>-Pho8Δ60 snf7Δ::nat* | this study |
| SSY2096 | ADE2 *pho13Δ::HIS3 pho8::kan-P<sub>GPD</sub>-Pho8Δ60 vps4Δ::nat* | this study |
| SSY2197 | ADE2 *pho8Δ::HIS3 pho13Δ::TRP1 YOP1-Pho8Δ60::kan nem1Δ::nat* | this study |
| SSY2346 | ADE2 *pho8Δ::HIS3 pho13Δ::TRP1 YOP1-Pho8Δ60::kan spo7Δ::nat* | this study |
| SSY2078 | ADE2 *pho8Δ::HIS3 pho13Δ::TRP1 YOP1-Pho8Δ60::kan vps23Δ::nat* | this study |
| SSY2377 | ADE2 *pho8Δ::HIS3 pho13Δ::TRP1 YOP1-Pho8Δ60::kan snf7Δ::nat* | this study |
| SSY2082 | ADE2 *pho8Δ::HIS3 pho13Δ::TRP1 YOP1-Pho8Δ60::kan vps4Δ::nat* | this study |
| SSY1194 | ADE2 *pho8Δ::HIS3 pho13Δ::TRP1 YOP1-Pho8Δ60::kan atg39Δ::nat* | this study |
| SSY1195 | ADE2 *pho8Δ::HIS3 pho13Δ::TRP1 YOP1-Pho8Δ60::kan atg40Δ::nat* | this study |
| SSY2175 | ADE2 *pho8Δ::HIS3 pho13Δ::TRP1 YOP1-Pho8Δ60::kan atg39Δ::nat atg40Δ::hph* | this study |
| SSY2425 | ADE2 *pho8Δ::HIS3 pho13Δ::TRP1 YOP1-Pho8Δ60::kan atg7Δ::LEU2 atg40Δ::nat* | this study |
| SSY2699 | ADE2 *leu2::P<sub>GAL1</sub>-GFP-Pho8-LEU2 (4x) Δvps23::hph* | this study |
| SSY2575 | ADE2 *leu2::P<sub>GAL1</sub>-GFP-Pho8-LEU2 (4x) Δsnf7::nat* | this study |
| SSY2574 | ADE2 *leu2::P<sub>GAL1</sub>-GFP-Pho8-LEU2 (4x) Δvps4::nat* | this study |
| SSY2430 | ADE2 *SEC63-mCherry::TRP1 leu2::P<sub>GAL1</sub>-gfp(Y66F)-Pho8-LEU2 (4x) his3::P<sub>VPS24</sub>-Snf7-LAP-mNeonGreen-URA3* | this study |
| SSY2801 | ADE2 *SEC63-mCherry::TRP1 leu2::P<sub>GAL1</sub>-gfp(Y66F)-Pho8-LEU2 (4x) his3::P<sub>SNF7</sub>-Snf7-LAP-mNeonGreen-URA3* | this study |
| SSY2705 | ADE2 *leu2::P<sub>GAL1</sub>-gfp(Y66F)-Pho8-LEU2 (4x) ura3::P<sub>VPS24</sub>-Snf7-LAP-mNeonGreen-URA3 trp1::P<sub>GPD</sub>-mCherry-Ubc6-TRP1* | this study |

