## Supplementary figures and images for "ESCRT machinery mediates selective microautophagy of endoplasmic reticulum"

### Movie 1.tif

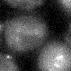

### Movie 2.tif

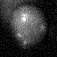
